# Niche-specific macrophage loss promotes skin capillary aging

**DOI:** 10.1101/2023.08.25.554832

**Authors:** Kailin R. Mesa, Kevin A. O’Connor, Charles Ng, Steven P. Salvatore, Alexandra Dolynuk, Michelle Rivera Lomeli, Dan R. Littman

## Abstract

All mammalian organs depend upon resident macrophage populations to coordinate repair processes and facilitate tissue-specific functions^1–3^. Recent work has established that functionally distinct macrophage populations reside in discrete tissue niches and are replenished through some combination of local proliferation and monocyte recruitment^4,5^. Moreover, decline in macrophage abundance and function in tissues has been shown to contribute to many age-associated pathologies, such as atherosclerosis, cancer, and neurodegeneration^6–8^. Despite these advances, the cellular mechanisms that coordinate macrophage organization and replenishment within an aging tissue niche remain largely unknown. Here we show that capillary-associated macrophages (CAMs) are selectively lost over time, which contributes to impaired vascular repair and tissue perfusion in older mice. To investigate resident macrophage behavior *in vivo*, we have employed intravital two-photon microscopy to non-invasively image in live mice the skin capillary plexus, a spatially well-defined model of niche aging that undergoes rarefication and functional decline with age. We find that CAMs are lost with age at a rate that outpaces that of capillary loss, leading to the progressive accumulation of capillary niches without an associated macrophage in both mice and humans. Phagocytic activity of CAMs was locally required to repair obstructed capillary blood flow, leaving macrophage-less niches selectively vulnerable to both homeostatic and injury-induced loss in blood flow. Our work demonstrates that homeostatic renewal of resident macrophages is not as finely tuned as has been previously suggested^9–11^. Specifically, we found that neighboring macrophages do not proliferate or reorganize sufficiently to maintain an optimal population across the skin capillary niche in the absence of additional cues from acute tissue damage or increased abundance of growth factors, such as colony stimulating factor 1 (CSF1). Such limitations in homeostatic renewal and organization of various niche-resident cell types are potentially early contributors to tissue aging, which may provide novel opportunities for future therapeutic interventions.

## Main Text

Tissue homeostasis is dependent on multiple macrophage populations that reside in distinct sub-tissue compartments or niches, such as epithelium, blood vessels or nerves, and are thought to support specialized tissue functions^4,12,13^. Recent work has suggested that functional decline and rarefication of vascular niches may contribute to various age-associated tissue pathologies (including sarcopenia, chronic wounds, and Alzheimer’s disease)^14–16^. It is not yet known how tissue-resident macrophages resist or potentiate such niche-specific aging processes^6,17^.

### Capillary-associated macrophage decline with age correlates with impaired blood flow

To model mammalian tissue aging, we adapted an intravital microscopy technique to visualize skin resident macrophage populations non-invasively in live mice throughout the lifetime of the organism^18,19^ (Figure 1a, Supplemental Videos 1 and 2). Unexpectedly, longitudinal imaging of skin macrophages (marked by *Csf1r-EGFP*) revealed niche-specific decline in macrophage populations. Subdividing the skin into three anatomical layers, epidermis, upper (papillary) and lower (reticulated) dermis, we observed that macrophages of the upper dermis were lost with age at a greater rate than macrophages from both epidermis and lower dermis (Figure 1b-c). Further characterization of this upper dermal CSF1R^+^ population revealed expression of additional macrophage markers^20–22^, including the chemokine receptor CX3CR1, lysozyme M (LysM), and the mannose receptor CD206 (Extended Data Fig. 1a,b,g,h). A major component of the upper dermal niche is the superficial capillary plexus, which supplies nutrient exchange for the overlying epidermis. To visualize this structure, we utilized third harmonic generation (3^rd^ Harmonic) from our imaging to track red blood cell (RBC) flow through capillary vessels^23,24^, which was comparable to conventional rhodamine dextran labeling (Extended Data Fig. 2 and Supplemental Video 3) and not significantly altered when utilizing a coverslip during our imaging sessions (Extended Data Fig. 3a,b). With this *in vivo* marker of RBC flow, we found that these macrophages were closely associated with blood capillaries of the superficial plexus, suggesting that they may provide support for this capillary niche (Extended Data Fig. 1c-e and Supplemental Video 4). To investigate if these macrophages play a role in capillary function, we assessed if RBC blood flow was altered in the presence of capillary-associated macrophages (CAMs). Performing timelapse recordings of fluorescently labeled CAMs in 2 month old mice with a cre-dependent dual reporter system (*Cx3cr1-CreERT2; R26-mTmG*), we found that capillaries lacking an associated macrophage had a higher rate of obstructed RBC flow (Figure 1d,e and Supplemental Video 5). We next performed longitudinal analysis across multiple ages (1-18 months) to assess CAM coverage and capillary function during physiological aging. We found across all ages tested a significant loss in RBC blood flow in capillaries lacking associated macrophages (Extended Data Fig. 3c). Furthermore, we found the fraction of capillaries with an associated macrophage significantly decreased with age (Figure 1f). This decrease in CAMs and coverage outpaced the loss of capillaries (Extended Data Fig. 3e,f), which was previously shown to be an early hallmark of mouse and human aging in multiple tissues, including central nervous system, lung, kidney, and skin^14,15,25–33^. Therefore, to assess if this phenomenon also occurs in humans, we obtained both young and old human patient skin samples. Consistent with our observations in mice, human capillary-associated macrophages also displayed a decline with age. Moreover, CAM decline also outpaced capillary loss with age, suggesting a similar loss in macrophage coverage of the capillary niche (Extended Data Fig. 3g-j). Therefore, these observations suggest that local macrophage loss in both mice and human may contribute to impaired capillary function with age. To assess any functional role of CAMs in maintaining capillary blood flow, we performed chemical and genetic ablation of CAMs, via clodronate liposomes and *Cx3cr1-DTR* depletion, respectively, and observed both acute CAM loss as well as loss in capillary flow (Figure 1 g,h and Extended Data Fig. 4). Collectively, this work highlights an evolutionarily conserved loss in skin capillary-associated macrophages with age, which correlates with impaired homeostatic capillary perfusion.

**Figure 1.**
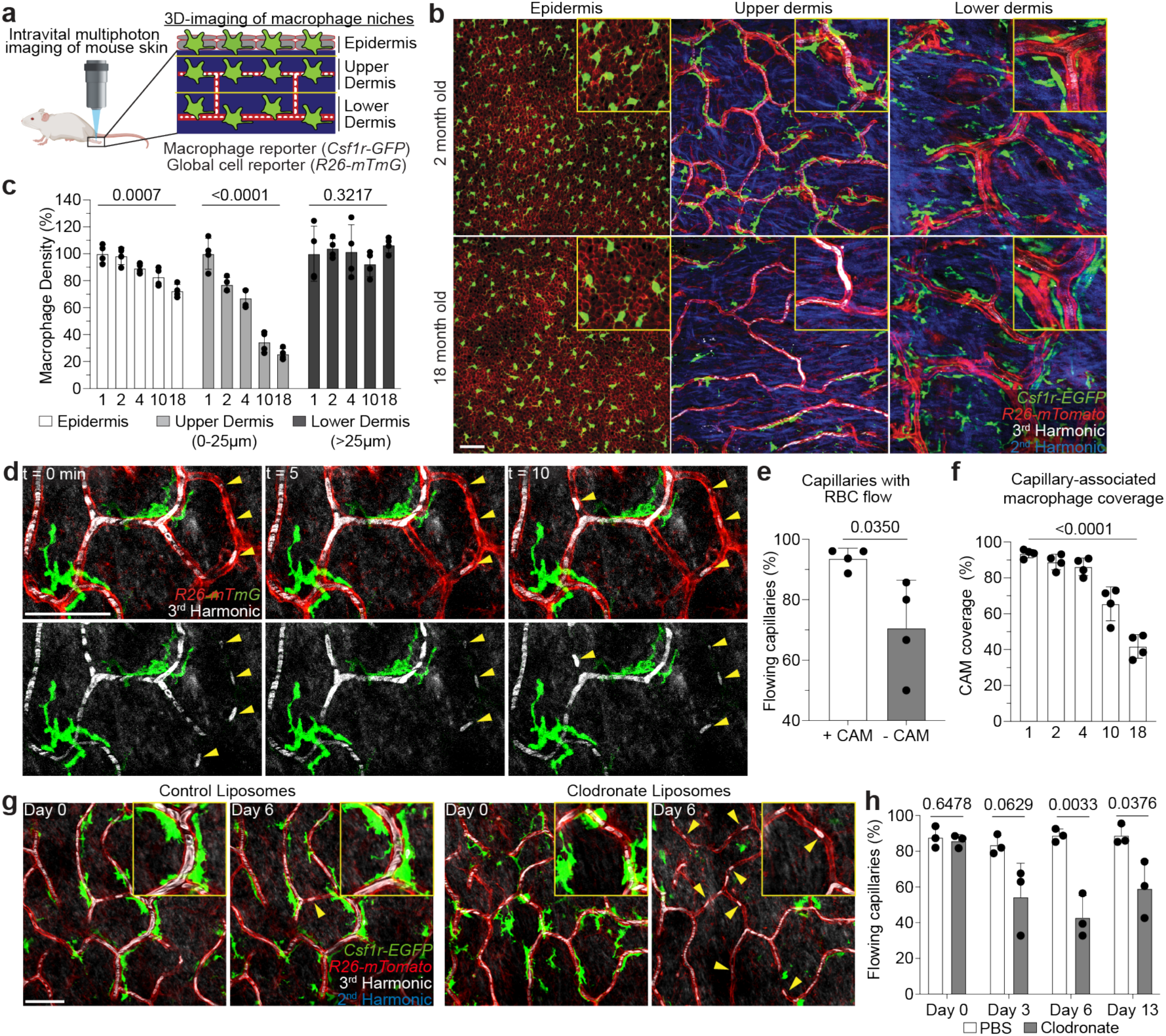
Niche-specific macrophage loss with age correlates with impaired skin capillary blood flow. **a,** Schematic of intravital imaging of resident macrophage populations in mouse skin, indicating epidermal and dermal populations, using the macrophage reporter, *Csf1r-EGFP,* in combination with an *Actb*-driven universal tdTomato reporter, *R26-mTmG*. **b,** Representative optical sections of distinct resident macrophage populations in young (2-month-old) and old (18-month-old) mice. **c,** Quantification of niche-specific macrophage density change in 1, 2, 4, 10, and 18 month old mice (n = 4 mice in each age group; two 500µm^2^ regions per mouse; macrophage density in each skin niche was compared across age groups by one-way ANOVA; mean ± SD). **d,** Skin resident macrophage labeling using *Cx3cr1-CreERT2;R26-mTmG* mice was performed following a single high-dose intraperitoneal injection of tamoxifen (2mg) in 1 month old mice. Single optical sections at successive time points 5 min apart showing red blood cell (RBC) flow (white) in capillaries (red) with or without nearby CAMs (green). Yellow arrowheads indicate obstructed RBC capillary flow. **e,** Quantification of capillaries with blood flow as measured by stalled RBCs as described in Extended Data Figure 2 (n = 226 CAM+ capillary segments, n = 27 CAM-capillary segments; n = 4 mice; capillary blood flow (CAM+ vs CAM-) was compared by paired Student’s t test; mean ± SD). **f,** Percentage of capillary segments with at least one associated macrophage (n = 4 mice in each age group; two 500µm^2^ regions per mouse; macrophage association with capillary segments was compared across age groups by one-way ANOVA; mean ± SD). **g,** Representative images demonstrate macrophage depletion following intradermal injections of clodronate-liposomes every 3 days. Repeated intravital imaging of the vascular niche was performed to visualize macrophages (*Csf1r-GFP*), capillaries (*R26-mTmG*) and RBC flow (Third Harmonic). **h,** Percentage of capillaries with blood flow following macrophage depletion (n = 194 capillary segments in clodronate group, n = 199 capillary segments in PBS group; 3 mice in each group; capillary flow (clodronate vs PBS) was compared by unpaired Student’s t test; mean ± SD). Scale bar, 50 µm.

### Local recruitment of CAMs is required to restore capillary blood flow

Given the link between CAMs and capillary blood flow, we next assessed the long-term fate of vessels that lack an associated macrophage. To this end, we performed a 6-month time course in *Cx3cr1-GFP*;*R26-mTmG* mice from 1 to 7 months of age (Figure 2a) and found that capillaries fated for pruning in the 6 month time course had decreased CAM coverage as compared to capillaries that were maintained (Figure 2b-c). This finding suggests that CAMs might be locally required to maintain proper capillary function and preservation with age. Therefore, to assess the cellular mechanism(s) by which CAMs support capillary function, we utilized a laser-induced blood clotting model to precisely target and stop blood flow in individual capillary segments (Figure 2d, Extended Data Fig. 5 and Supplemental Video 6). To assess macrophage involvement, we tracked the daily displacement of surrounding CAMs to laser-induced clots. These data showed that CAM recruitment to sites of capillary damage as well as RBC engulfment are locally restricted to within approximately 80µm, and largely occur within the first two days after injury (Figure 2e,f). This is consistent with previous work describing macrophage cloaking as an acute behavioral response to laser-induced tissue damage^34,35^. Multiple signaling pathways are involved in sensing tissue damage^34,36,37^, including the chemokine receptor CX3CR1, which is homeostatically expressed by CAMs (Extended Data Fig. 1a-f). Therefore, we interrogated the role of CX3CR1 signaling in recruiting nearby CAMs to capillary damage. We performed laser-induced clotting and found significant impairment in CAM recruitment in *Cx3cr1-gfp/gfp* mice as compared to *Cx3cr1-gfp/+* controls (Figure 2g-h). We also found that CX3CR1-deficiency led to a significant delay and overall reduction in capillary reperfusion by Day 7 post clot induction (Figure 2i). From these findings, we assessed capillary RBC flow during physiological aging and found a significant loss in flowing capillaries in *Cx3cr1-gfp/gfp* mice, as compared to *Cx3cr1-gfp/+* controls, by 6 months of age (Figure 2j). Together, our results support a model where local CAM recruitment, in part through CX3CR1-signaling, is critical for both capillary repair and the long-term preservation of the vascular network during physiological aging.

**Figure 2.**
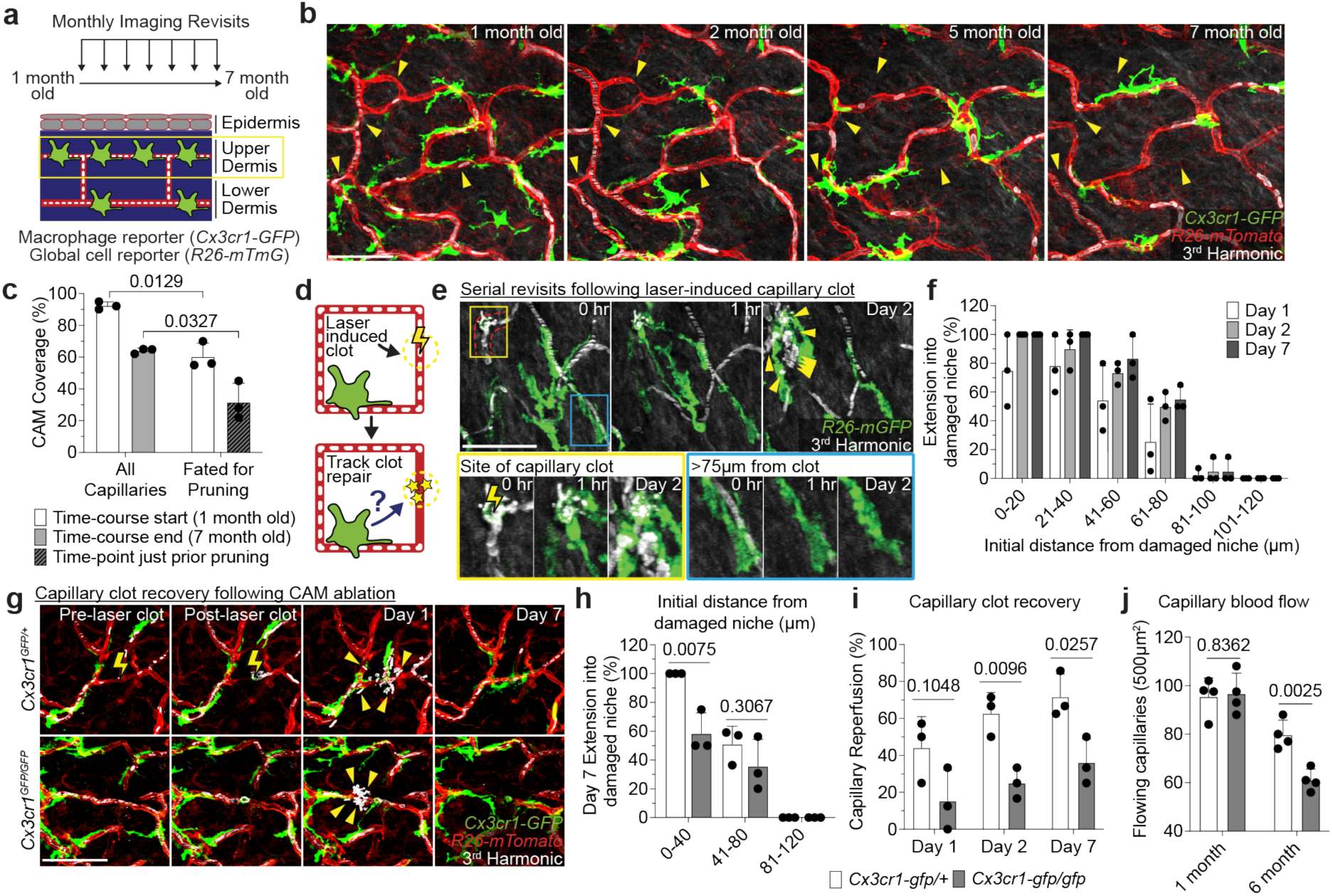
Local recruitment of CAMs is required to restore capillary blood flow. **a,** Scheme of long-term serial imaging of capillary niche in *Cx3cr1-GFP;R26-mTmG* mice. **b,** Sequential revisits reveal progressive pruning of capillary niche over a 6-month period. Yellow arrowheads indicate capillaries that will undergo pruning. **c,** Quantification of capillary-associated macrophage coverage at 1 and 7 months of age (n = 3 mice; three 500µm^2^ regions per mouse; Frequency of macrophage association with capillary segments was compared between capillaries fated for pruning and all capillaries at 1-and 7-month-old time-points by paired Student’s t test; mean ± SD). **d,** Scheme of laser-induced capillary clot experiment. **e,** Sequential revisits of damaged capillary segment (red dashed lines) after laser-induced clot formation in *Cx3cr1-CreERT2;R26-mTmG* mice. Yellow lightning bolt indicates site of laser-induced capillary clot. Yellow arrowheads indicate extra-luminal vascular debris. **f,** Quantification of capillary-associated macrophage extension toward damaged niche at Day 1, 2 and 7 after laser-induced clotting (n = 50 CAMs, in 3 mice; mean ± SD). **g,** Sequential revisits of damaged capillary niche after laser-induced clot formation (yellow lightning bolt) in *Cx3cr^gfp/+^* and *Cx3cr^gfp/gfp^* mice. CAMs (green), capillaries (red), RBC (white). Yellow arrowheads indicate extra-luminal vascular debris. **h,** Quantification of capillary-associated macrophage extension toward damaged niche at Day 7 after laser-induced clotting (n = 99 CAMs in *Cx3cr^gfp/+^* group; n = 65 CAMs in *Cx3cr^gfp/gfp^* group; n = 3 mice in each group; CAM extension (*Cx3cr^gfp/+^*vs *Cx3cr^gfp/gfp^*) was compared by unpaired Student’s t test; mean ± SD). **i,** Quantification of capillary reperfusion at Day 1, 2 and 7 after laser-induced clotting (n = 20 clots in *Cx3cr^gfp/+^*group; n = 21 clots in *Cx3cr^gfp/gfp^* group; n = 3 mice in each group; capillary reperfusion (*Cx3cr^gfp/+^* vs *Cx3cr^gfp/gfp^*) was compared by unpaired Student’s t test; mean ± SD). **j,** Quantification of capillary blood flow (n = 4 mice in each group; two 500µm^2^ regions per mouse; capillary blood flow (*Cx3cr^gfp/+^* vs *Cx3cr^gfp/gfp^*) was compared at 1 and 6 months of age by unpaired Student’s t test; mean ± SD). Scale bar, 50µm.

### CAMs are required to clear vascular damage and preserve skin capillaries during aging

To determine if diminished capillary function with age is directly related to macrophage loss, we took advantage of the inherent variability in CAM density in older mice. Specifically, we performed the laser-induced clotting of aged skin capillaries with or without local CAMs (within 75µm from clot) (Figure 3a). Remarkably, capillaries that retained local CAMs in aged mice were significantly better at reestablishing blood flow as compared to capillaries without local CAMs in the same mice (Figure 3b). This result suggests that local macrophage loss drives age-associated capillary dysfunction. To directly test the role of macrophages in capillary repair, we performed laser-induced ablation of local CAMs immediately prior to capillary clot-induction. Indeed, in regions where CAMs were ablated, repair was significantly impaired and blood flow was not properly reestablished (Figure 3c,d). We targeted other perivascular cells with the same laser ablation conditions and found no impairment to capillary repair, nor did we detect neutrophil swarming as has been described for much larger areas of laser-induced damage^34,35,38–44^ (Extended Data Fig. 6 and 7).

**Figure 3.**
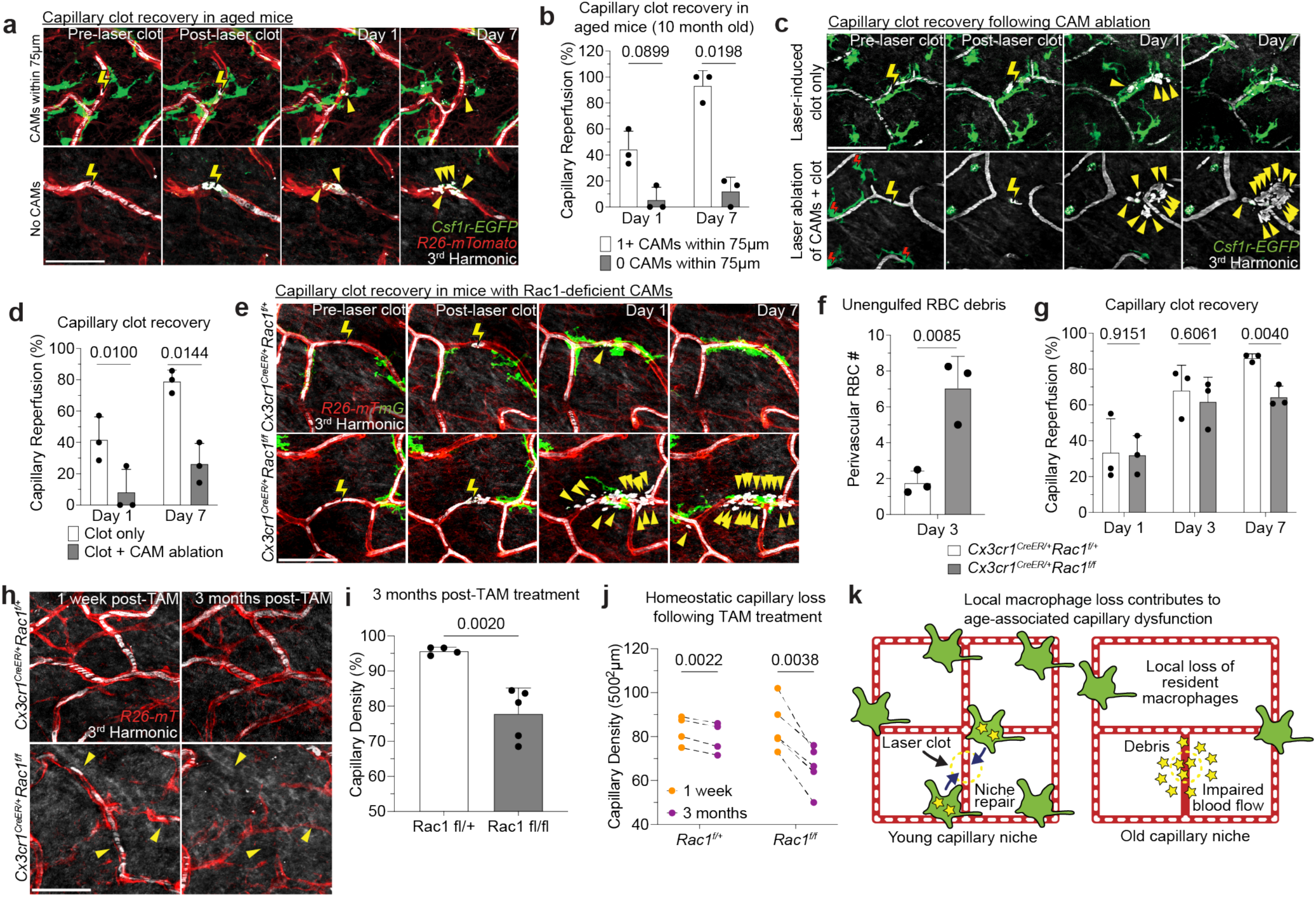
CAMs are required to clear vascular damage and preserve skin capillaries during aging. **a,** Sequential revisits of damaged capillary niche after laser-induced clot formation in *Csf1r-EGFP;R26-mTmG* mice. Capillary clot formation (yellow lightning bolt) was performed at 940nm for 1s in 10-month-old mice. Yellow arrowheads indicate extra-luminal vascular debris. **b,** Quantification of capillary reperfusion at day 1 and 7 after laser-induced clotting (n = 16 capillary clots in regions with CAMs (<75µm from clot), n = 16 capillary clots in regions without CAMs (>75µm from clot; 3 mice in total; capillary reperfusion (with CAMs vs without CAMs) was compared at Day 1 and Day 7 by paired Student’s t test; mean ± SD). Scale bar, 50 µm. **c,** Sequential revisits of damaged capillary niche after laser-induced CAM ablation and clot formation in *Csf1r-EGFP* mice. Macrophage laser ablation (red lightning bolt) and capillary clot formation (yellow lightning bolt) were both performed at 940nm for 1s. **d,** Quantification of capillary reperfusion at Day 1 and 7 after laser-induced clotting and macrophage ablation (n = 19 capillary clots in CAM ablated regions, n = 16 capillary clots in control regions; 3 mice in total; capillary reperfusion (CAM ablated vs control) was compared by paired Student’s t test; mean ± SD). **e,** Sequential revisits of damaged capillary niche after laser-induced clot formation in *Cx3cr1^CreER^*;*Rac1^fl/fl^* and *Cx3cr1^CreER^*;*Rac1^fl/+^*mice. CAMs (green), capillaries (red), RBC (white). **f,** Quantification of perivascular red blood cell debris at Day 3 after laser-induced clotting (n = 52 clots in *Rac1^fl/+^* group; n = 48 clots in *Rac1^fl/fl^*group; n = 3 mice in each group; capillary reperfusion (*Rac1^fl/+^*vs *Rac1^fl/fl^*) was compared by unpaired Student’s t test; mean ± SD). **g,** Quantification of capillary reperfusion at Day 1, 3 and 7 after laser-induced clotting (n = 67 clots in *Rac1^fl/+^* group; n = 82 clots in *Rac1^fl/fl^* group; n = 3 mice in each group; capillary reperfusion (*Rac1^fl/+^* vs *Rac1^fl/fl^*) was compared by unpaired Student’s t test; mean ± SD). **h,** Sequential revisits reveal pruning of capillaries over a 3-month period. Yellow arrowheads indicate capillaries that will undergo pruning. **i,** Quantification of capillary density (n = 4 mice in *Rac1^fl/^* group and 5 mice in *Rac1^fl/fl^* group; two 500µm^2^ regions per mouse; capillary density (*Rac1^fl/+^* vs *Rac1^fl/fl^*) was compared at 3 months post tamoxifen treatment by unpaired Student’s t test; mean ± SD). **j,** Quantification of capillary density (n = 4 mice in *Rac1^fl/^* group and 5 mice in *Rac1^fl/fl^*group; two 500µm^2^ regions per mouse; capillary density (1 week vs 3 months) was compared in *Rac1^fl/+^* and *Rac1^fl/fl^*mice by paired Student’s t test; mean ± SD). **k,** Scheme of age-associated decline of skin capillary niche. Capillary-associated macrophage density declines with age, which predisposes aged capillaries to impaired repair and sustained tissue perfusion. Scale bar, 50µm.

From our serial imaging of capillary repair, we found nearby CAMs often surrounded by and containing RBC debris. To mechanistically understand if CAM uptake and clearance of this vascular debris is functionally important for capillary repair, we acutely impaired CAM phagocytosis through inducible cre-dependent knockout of Rac1, a critical component of the phagocytic machinery^45^, one week prior to laser-induced clot formation. While there was no significant change in CAM density (Extended Data Fig. 8), there was significant impairment in clearance of RBC debris and capillary reperfusion in *Cx3cr1^CreER^*;*Rac1^fl/fl^* mice compared to *Cx3cr1^CreER^*;*Rac1^fl/+^*littermate controls (Figure 3e-g), suggesting that Rac1-dependent phagocytic clearance is critical for proper capillary repair and tissue reperfusion.

To test the long-term vascular effects of Rac1-deficiency in CAMs, we assessed capillary density three months after tamoxifen administration in *Cx3cr1^CreER^*;*Rac1^fl/fl^*mice. We found an acceleration in rate of capillary pruning after three months of physiological aging in the skin of *Cx3cr1^CreER^*;*Rac1^fl/fl^* mice compared to *Cx3cr1^CreER^*;*Rac1^fl/+^*littermate controls (Figure 3h-j). Therefore, it is likely that loss of Rac-1 dependent behaviors, such as phagocytosis of vascular debris, directly contributes to impaired recovery and preservation of the skin microvascular network.

Together, our results support a model in which local CAM recruitment and phagocytic clearance of vascular debris is critical to maintain capillary function. Therefore, as CAM density declines with age, so does capillary perfusion of the tissue (Figure 3k).

### Dermal macrophages utilize niche-specific self-renewal strategies leading to selective CAM loss with age

Maintenance of tissue resident macrophage populations in the skin is thought to be mediated through a combination of local proliferation and systemic replacement by bone marrow-derived blood monocytes^13,46,47^. To understand how CAMs are replenished under physiological conditions, we generated bone marrow chimeras with *Csf1r-GFP*;*CAG-dsRed* bone marrow transferred into lethally irradiated *Csf1r-GFP* mice. Importantly, the hind paws of these mice were lead-shielded to prevent loss of resident macrophage populations from our imaging area (Figure 4a). Tracking macrophage populations from all anatomical layers of the skin for 10 weeks post transplantation showed that while most lower dermal macrophages are replaced by monocytes, <5% of upper dermal macrophages were bone marrow-derived (Figure 4b,c). Additionally, capillaries with either bone marrow-derived (GFP+/dsRed+) or host-derived (GFP+/dsRed-) CAMs showed no obvious differences in RBC blood flow (Extended Data Fig. 3d), suggesting that CAM ontogeny may not play a major role in homeostatic capillary function.

**Figure 4.**
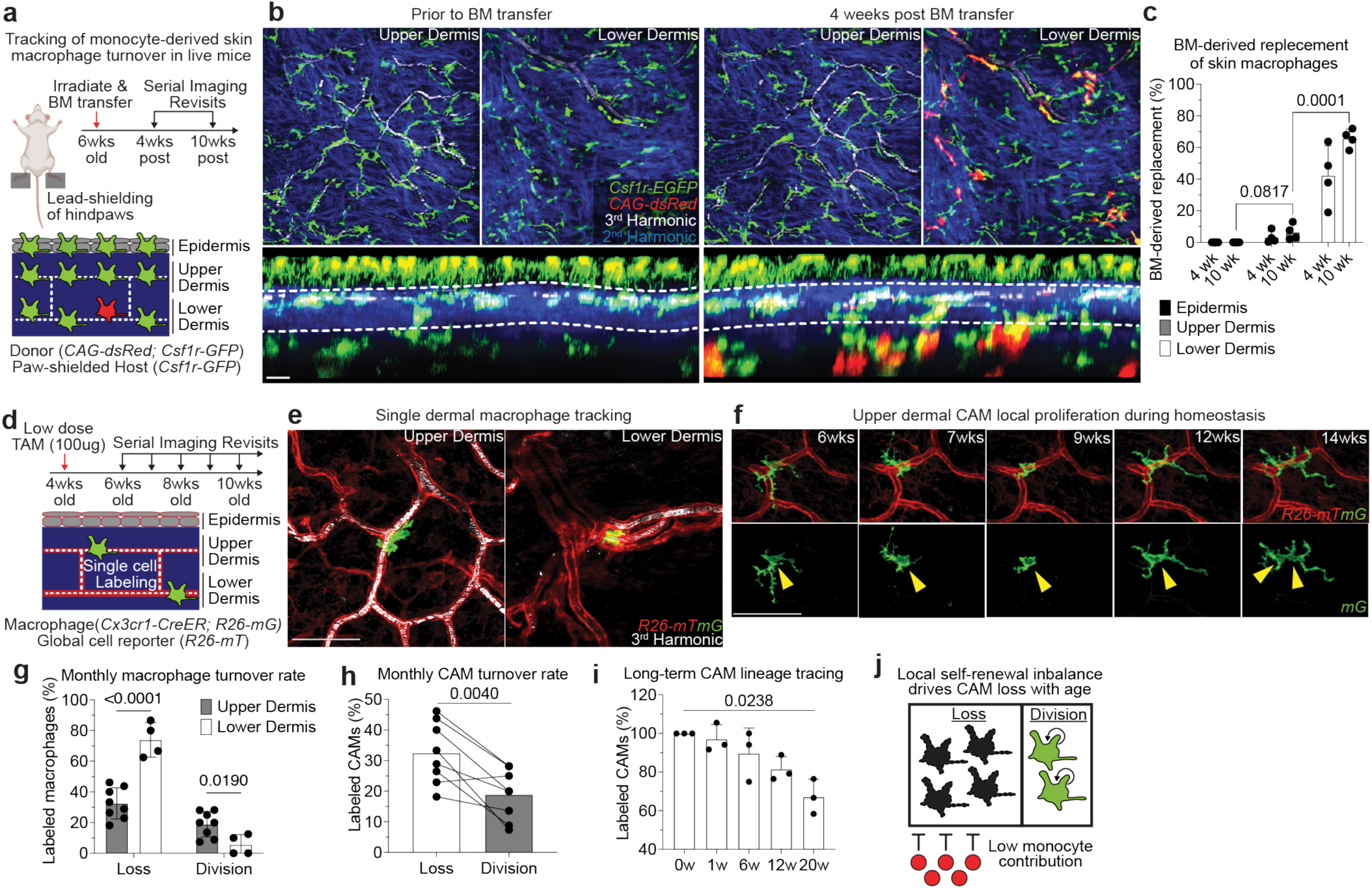
Dermal macrophages utilize niche-specific self-renewal strategies leading to selective CAM loss with age. **a,** Scheme of serial imaging of skin resident macrophage repopulation in bone marrow chimeras with *Csf1r-GFP*;*CAG-dsRed* bone marrow transferred into lethally irradiated *Csf1r-GFP* mice. Note hind paws of these mice were lead-shielded to prevent irradiation-induced loss of resident macrophage populations from our imaging area. **b,** Representative optical sections of dermal macrophage populations in mice prior to BM transfer and 4 weeks post BM transfer. **c,** Quantification of macrophage repopulation at 4-and 10-weeks post bone marrow transplantation (n = 4 mice; two 500µm^2^ regions of each skin compartment per mouse; bone marrow-derived macrophage replacement (percentage of GFP+/dsRed+ macrophages) was compared between skin compartments at 10 weeks post BM transplantation by paired Student’s t test; mean ± SD). **d,** Scheme of long-term tracking of dermal macrophage self-renewal experiment. **e,** Representative images in upper and lower dermis of single macrophage tracing in *Cx3cr1-CreERT2*; *R26-mTmG* mice. Weekly revisits were performed during homeostatic conditions for 20 weeks following a single low-dose intraperitoneal injection of tamoxifen (50µg). **f,** Representative serial revisit images of single macrophage tracing in *Cx3cr1-CreERT2*; *R26-mTmG* mice. **g,** Quantification of monthly rate of dermal macrophage loss and division in upper (n = 262 macrophages; 8 mice in total) and lower (n = 42 macrophages; 4 mice in total) dermis; macrophage monthly division and loss rates (upper vs lower dermis) were compared by unpaired Student’s t test; mean ± SD. **h,i,** Quantification of (h) monthly rate of CAM loss and division (n = 262 CAMs; 8 mice in total; CAM turnover rates (CAM loss vs division) was compared by paired Student’s t test; mean ± SD) and of (i) maintenance of the labeled CAM population over 20wk lineage tracing (n = 59 CAMs; 3 mice in total; number of fluorescently-labeled CAMs was compared across each time point in the same mice by one-way repeated measures ANOVA with Geisser-Greenhouse correction; mean ± SD). **j,** Scheme of local self-renewal imbalance drives CAM loss with age. In contrast to lower dermal macrophages, which are continuously replenished by monocytes, CAMs of the upper dermis largely rely of local proliferation. However, the rate of CAM division is insufficient to stably maintain this population with age. Scale bar, 50µm.

Given that nearly all CAMs remained host-derived, we next performed single-macrophage lineage tracing to monitor local proliferation of dermal macrophages. Specifically, we induced sparse cre-recombination in 1 month old *Cx3cr1-GFP*;*R26-mTmG* mice to label and track individual macrophages from both the upper and lower dermis over weekly revisits (Figure 4d-e). From these serial revisits, we observed striking positional stability of macrophages on the vascular network as well as local cell division (Figure 4f). Analysis of monthly rates of macrophage loss and division revealed that lower dermal macrophages have a significantly lower division rate as well as significantly higher loss rate, as compared to CAMs (upper dermal macrophages) (Figure 4g).

Focusing on CAM turnover, we also found a significant skew toward cell loss over division during the 4-month time course (Figure 4h). Consistent with these data, there was a significant decline in the fraction of fate-mapped CAMs over the 20-week time course (Figure 4i). These results demonstrate that CX3CR1+ dermal macrophage self-renewal strategies are niche-specific, where blood monocyte recruitment is utilized in the lower dermis and local cell division is utilized in the upper dermis. Importantly, we find that CAMs from the upper dermis are insufficiently replenished by local proliferation, which contributes to their progressive decline in this tissue niche (Figure 4j).

### Macrophage loss without local tissue damage is not sufficient to promote CAM renewal

The ability of resident macrophages to locally self-renew has largely been studied through methods of near-total macrophage depletion, which have limited niche-specificity and often generate tissue-wide inflammation^9–11^. To directly interrogate the steps of macrophage self-renewal over time, we tracked both the replacement of individual macrophages after loss and the redistribution of sister macrophages after division. First, to track local macrophage replacement after CAM loss, we performed laser-induced ablation of all CAMs within a defined 500µm^2^ region. Serial revisits up to two weeks after ablation revealed minimal repopulation from adjacent capillary regions that retained intact CAM populations (Figure 5a). We found a similar lack in repopulation following partial CAM depletion in *Cx3cr1^DTR^* mice following low dose diphtheria toxin administration (Figure 5b). To avoid any non-physiological effects from these cell depletion models, we developed a dual fluorescent macrophage reporter mouse, *Cx3cr1-CreERT2; Rosa26-dsRed; Csf1r-EGFP*, allowing for cre-dependent recombination to differentially label a small fraction of macrophages and track their homeostatic replacement. Examination on serial weekly revisits indicated that, as we observed in depletion models, most capillary niches did not recruit a new macrophage for as long as two weeks following CAM loss (Figure 5c,d). In contrast, when we employed large laser-induced damage (500µm^2^ region) in either the upper dermis or overlaying epidermis (Figure 5e and Extended Data Fig. 9a) we found CAMs were readily replenished (Figure 5e-g), in a CCR2-dependent manner, suggesting at least partial monocyte repopulation of CAMs after tissue damage. We also found that local CAM proliferation was significantly increased one week following laser-induced damage in either the upper dermis or epidermis (Figure 5h,i and Extended Data Fig. 9a-c). Taken together, these results suggest that CAM loss alone is not a sufficient trigger to promote macrophage replenishment and requires additional signals elicited by tissue damage.

**Figure 5.**
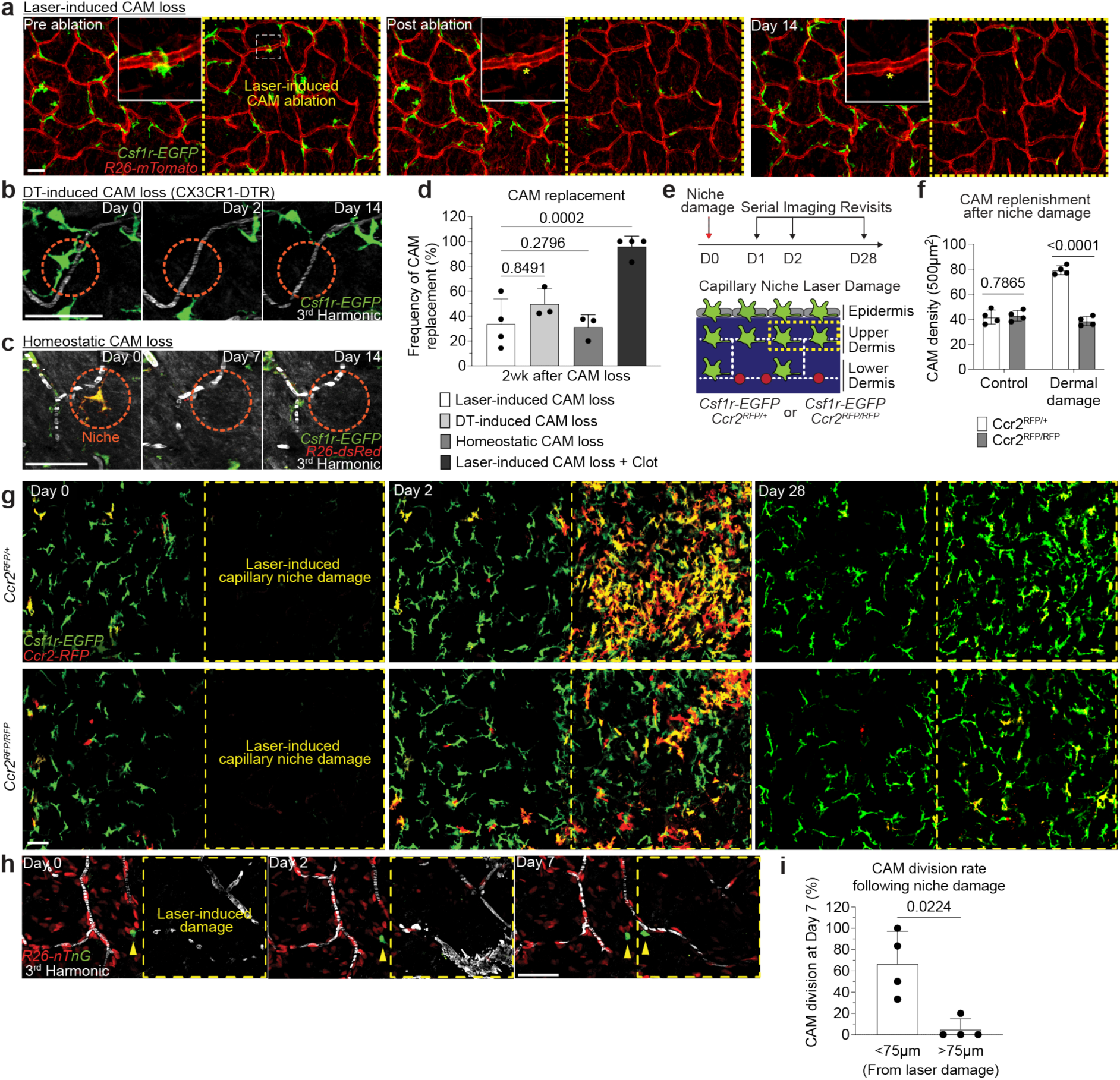
Macrophage loss without local tissue damage is not sufficient to promote CAM renewal. **a,** Representative revisits of CAM replacement after laser-induced CAM ablation in *Csf1r-EGFP;R26-mTmG* mice. Macrophage laser ablations (yellow asterisk) were performed in a 500µm^2^ region (yellow dashed line square), **b,** Representative revisits of CAM replacement after a single intraperitoneal injection of diphtheria toxin (25ng/g body weight in PBS) in *Cx3cr1-DTR* mice. **c,** Representative revisits of single lineage-tracked macrophages in *Cx3cr1-CreERT2*; *R26-dsRed*; *Csf1r-EGFP* mice. Weekly revisits were performed during homeostatic conditions for 2 weeks following a single low-dose intraperitoneal injection of tamoxifen (50µg). **d,** Quantification of replenishment rate of empty CAM niche under laser-induced loss (n = 44 CAMs; 4 mice in total), DT-induced loss (n = 53 CAMs; 3 mice in total), homeostatic loss (n = 29 CAMs; 3 mice in total), or laser-induced CAM loss with capillary clot (n = 25 CAMs; 4 mice in total). Frequency of CAM replacement after forced and homeostatic CAM loss were compared by unpaired Student’s t test; mean ± SD. **e,** Scheme of CAM replacement after capillary niche damage in *Csf1r-EGFP*; *Ccr2^RFP/+^*or *Ccr2^RFP/RFP^* mice. **f,** Quantification of replenishment rate (*Csf1r-EGFP*+ cells) of empty CAM niche after laser-induced CAM loss and niche damage (n = 4 mice per group; two 500µm^2^ regions per mouse; CAM density (*Ccr2^RFP/+^* vs *Ccr2^RFP/RFP^*) was compared by unpaired Student’s t test; mean ± SD). **g,** Representative revisits of CAM replacement after a regional laser-induced damage to the capillary niche. **h,** Representative revisits of single macrophage lineage tracing in *Cx3cr1-CreERT2*; *R26-nTnG* mice following capillary damage. **i,** Quantification of CAM proliferation based on proximity to capillary damage (n = 25 CAMs tracked in capillary damage regions, 4 mice in each group; CAM proliferation (based on damage proximity) was compared at day 7 by paired Student’s t test; mean ± SD). Scale bar, 50µm.

Second, to precisely track if proliferating CAMs readily redistribute across the capillary network following cell division, we utilized mice with a dual fluorescent nuclear reporter, *Cx3cr1-CreERT2; R26-nTnG*, and administered a low or high dose of tamoxifen to label either a sparse subset or all CAMs, respectively (Extended Data Fig. 9d). Compared to the average distance between all nearest neighboring macrophages, sister CAMs remained significantly closer (<15µm) to each other for at least 2 weeks after division (Extended Data Fig. 9e,f). These findings further support the notion that CAM division and loss are not spatiotemporally coupled, which we predict would progressively lead to disorganized patterning and the accumulation of both empty and crowded capillary regions. We tested this prediction by looking at the distribution of neighboring CAMs in both young (2-month-old) and old (10-month-old) mice. In young mice, the majority of macrophages were within 50µm of each other. In contrast, old mice had a biphasic distribution of macrophage patterning, with most CAMs either within 25µm or further than 75µm apart (Extended Data Fig. 9g). These results highlight two distinct cellular features that contribute to reduced CAM coverage with age: 1) insufficient macrophage repopulation following CAM loss, and 2) insufficient redistribution of these cells along the capillary niche, which may promote progressive erosion of the vascular network (Extended Data Fig. 9h).

### Local CAM expansion in old mice is sufficient to rejuvenate capillary repair and tissue reperfusion

While homeostatic CAM proliferation was insufficient to maintain a stable population density, we asked if extrinsic cues such as large tissue damage could increase CAM density long-term to improve capillary function in older mice. To this end, we found that large laser-induced epidermal damage (500µm^2^ region) resulted in a lasting increase in CAMs below the damaged regions compared to neighboring control regions (Extended Data Fig. 9i-k). Furthermore, we found a significant improvement in capillary repair in these same regions following laser-induced clotting (Extended Data Fig. 9l).

Our results suggest that CAMs in old mice can be stably expanded following environmental changes, such as tissue damage. Therefore, we next assessed if directly increasing CAM density, without local damage, would also be sufficient to improve future capillary repair and reperfusion. To this end, we utilized a fusion protein of the canonical macrophage growth factor, CSF1, with the Fc region of porcine IgG (CSF1-Fc), as it has been reported to robustly increase macrophage density in multiple tissues, including skin^48–50^. We performed daily intradermal injections of either CSF1-Fc or PBS in the left or right hind paws, respectively, of the same mice (Figure 6a). There was a significant increase in CAMs in the CSF1-treated paws compared to contralateral PBS controls, which had no significant change from pretreatment (Figure 6b,c).

**Figure 6.**
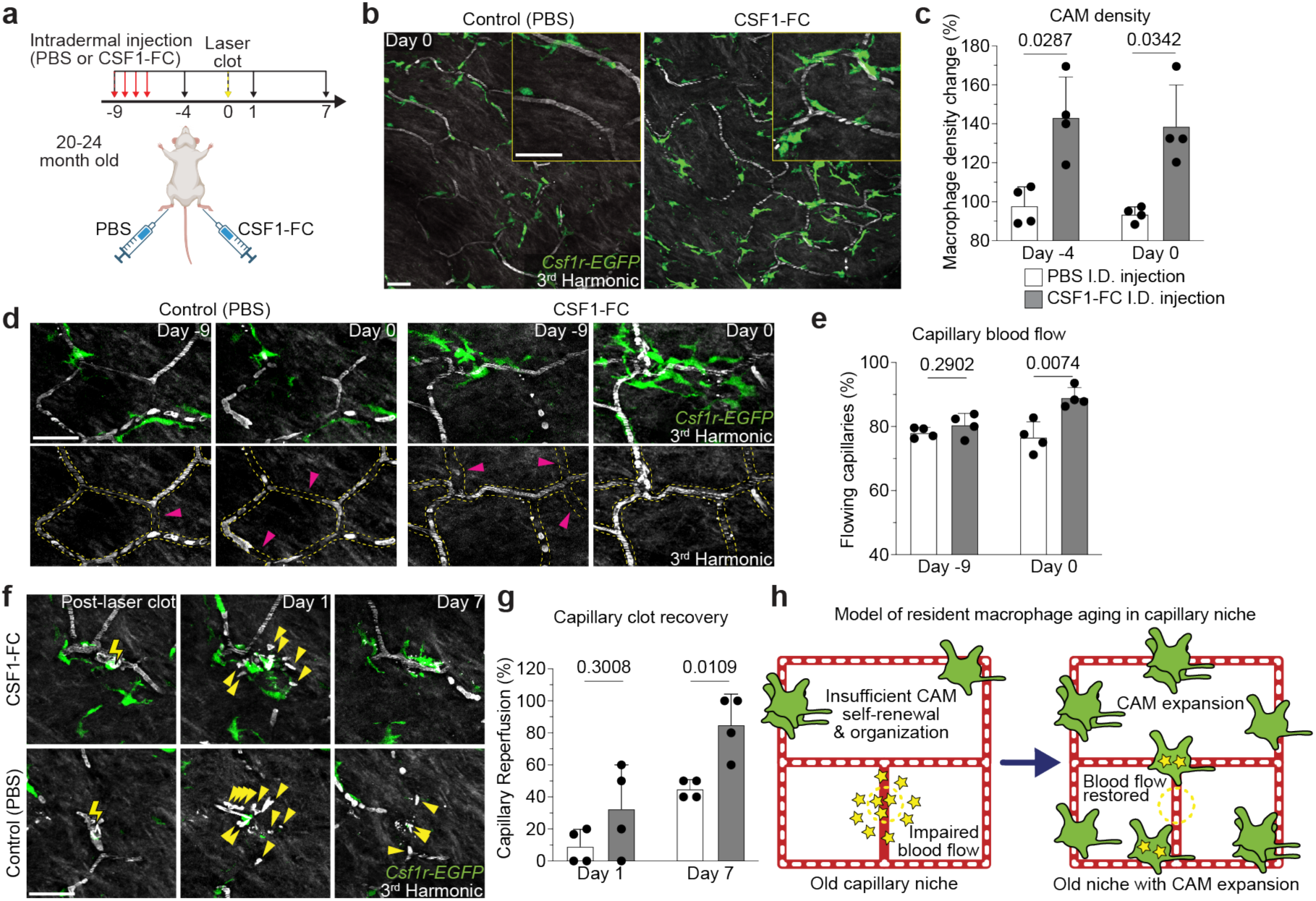
Local CAM replenishment in old mice is sufficient to rejuvenate capillary repair and tissue reperfusion. **a,** Scheme of CSF1-induced rejuvenation of aged capillary niche. **b,** Representative images of CAM density in *Csf1r-EGFP* mice nine days following daily intradermal injections (4 days) of CSF1-Fc (porcine CSF1 and IgG1a Fc region fusion protein) and PBS in contralateral hind paws of 20–24-month-old mice. **c,** Quantification of CAM density following CSF1-Fc or PBS treatment (n = 4 mice in total; two 500µm^2^ regions of each treatment condition per mouse; Percentage of CAM density change from Day -9 (CSF1-Fc vs PBS) was compared for Day -4 and Day 0 time points by paired Student’s t test; mean ± SD). **d,** Representative images of capillary (red dashed lines) blood flow in *Csf1r-EGFP* mice nine days following daily intradermal injections of CSF1-Fc and PBS in contralateral hind paws of 20–24-month-old mice. Magenta arrowheads indicate obstructed RBC capillary flow. **e,** Quantification of capillary blood flow following CSF1-Fc or PBS treatment (n = 214 capillary segments in CSF1-Fc treated regions (grey bar graph), n = 214 capillary segments in PBS treated regions (white bar graph); n = 4 mice in total; capillary blood flow (CSF1-Fc vs PBS) was compared for Day -9 and Day 0 time points by paired Student’s t test; mean ± SD). **f,** Sequential revisits of damaged capillary niche after laser-induced clot formation (yellow lightning bolt) in *Csf1r-EGFP* mice. Yellow arrowheads indicate extra-luminal vascular debris. **g,** Quantification of capillary reperfusion at Day 1 and 7 after laser-induced clotting (n = 18 clots in CSF1-Fc group (grey bar graph); n = 20 clots in PBS group (white bar graph); n = 4 mice in total; capillary reperfusion (CSF1-Fc vs PBS) was compared by paired Student’s t test; mean ± SD). **h,** Working model of resident macrophage aging in the skin capillary niche. Age-associated impairment in capillary repair can be rejuvenated following local expansion of resident macrophage population. Scale bar, 50µm.

To assess if CSF1-treatment modulated local CAM survival/proliferation or simply recruited new bone marrow-derived macrophages from blood monocytes, we performed CSF1-treatment in chimeric bone marrow mice where we could track the relative expansion of local and recruited populations (Extended Data Fig. 10a,b). Consistent with our previous experiments, we found that PBS injected paws showed no significant increase in host-or BM-derived macrophages in our imaging areas (Extended Data Fig. 10c-d). Importantly we found that CSF1-induced CAM expansion did not alter the ratio of resident (GFP+) and recruited (GFP+/dsRed+) CAMs (Extended Data Fig. 10e), as a relative increase in GFP+/dsRed+ CAMs over GFP+ CAMs would have suggested BM-derived monocyte recruitment. Therefore, this strongly suggests that CSF1 drives local proliferation of the existing CAM population.

Strikingly, we found that CSF1-treatment was sufficient to improve homeostatic capillary blood flow in old mice, as compared to PBS controls which had significantly more obstructed capillary segments (Figure 6d,e). Utilizing the same aged mice, we next tested if this increase in CAM density would be sufficient to improve capillary repair rates. Following laser-induced clotting, there was a significant improvement in capillary repair and reperfusion in CSF1-treated mice compared to PBS controls (Figure 6f,g), demonstrating that restoring dermal macrophage density in old mice can improve age-associated vascular dysfunction.

## Discussion

Macrophage renewal has largely been studied in non-physiological settings, such as through *in vitro* cell culture or severe depletion models that often are accompanied by acute inflammation^9,10,51–53^. Our work clearly demonstrates that the homeostatic renewal strategies of resident macrophages is niche-specific and not as finely tuned as has been previously suggested. Specifically, we found that macrophages of the upper dermis do not proliferate or redistribute sufficiently to maintain an optimal coverage across the skin capillary network unless they receive additional cues from acute tissue damage or increased growth factor abundance (Figure 6h). Interestingly, we confirmed previous findings that show epidermal Langerhans cell density also declines with age, which has been associated with impaired epidermal function^54–57^. This raises the possibility that age-associated loss in macrophage density is a more general phenomenon in populations that rely on local self-renewal.

In addition to self-renewal, we also found that CAM recruitment to repair tissue damage was spatially restricted. To our knowledge, the long-term size and stability of resident macrophage territories or niches *in vivo* has not been reported. Additionally, our work provides strong evidence that injury-induced dermal macrophage recruitment is restricted to approximately 80µm, as has been demonstrated in other tissues^34,35^. In young mice, CAM density is high enough to provide substantial niche/territory overlap between neighbors. However, with declining CAM density with age, we show that a significant fraction of the skin capillary network is no longer within a CAM’s territory range. It will be important to understand how this property of resident macrophages is influenced by other aspects of regional heterogeneity, such as innate immune imprinting^13^, to shape local immune responses in tissues.

Collectively, this work demonstrates that loss in CAMs: 1) begins within the first few months of life, 2) is progressive throughout life, and 3) is functionally detrimental to vascular function and preservation, which has been shown to be a primary driver of age-associated tissue impairments^15,58^. Furthermore, this work provides a novel platform to investigate age-associated deviations in tissue homeostasis at the single-cell level in a living mammal.

## Supporting information

Supplemental Video 1. Serial optical sections through mouse plantar skin. All cells (red), dermal collagen (blue) and red blood cells (white).

Supplemental Video 2. Serial optical sections of mouse skin macrophage populations (green) and dermal collagen (blue).

Supplemental Video 3. Time-lapse recording of capillary blood flow via red blood cells (white) and intravenous dextran (red).

Supplemental Video 4. Serial optical sections through upper dermal capillary niche in Cx3cr1-GFP; R26-mTmG mice.

Supplemental Video 5. Time-lapse recording of inconsistent and obstructed blood flow (white) in capillary (red) segments without macrophages (green).

Supplemental Video 6. Time-lapse recording following laser-induced capillary clot formation with rapid migration of CAMs (green) toward site of clot.

## Acknowledgements

We thank members of the Littman lab for valuable discussion and critical reading of the manuscript. We thank Michael Cammer from the NYU Microscopy Core for valuable discussion, training and technical support. The Microscopy Core is partially supported by NYU Cancer Center Support Grant NIH/NCI P30CA016087 at the Laura and Isaac Perlmutter Cancer Center, S10 RR023704-01A1 and NIH S10 ODO019974-01A1. This work was supported by a Jane Coffin Childs Fund fellowship (K.R.M.), Kirschstein-NRSA training grant T32AR64184 (K.R.M.), Kirschstein-NRSA individual postdoctoral fellowship F32AG071336 (K.R.M.), a Charles H. Revson Senior Fellowship in Biomedical Science (K.R.M.), the Helen and Martin Kimmel Center for Biology and Medicine (D.R.L.), NIH grant R01AI158687 (D.R.L.), and the Howard Hughes Medical Institute (D.R.L.).

## Author Contributions

K.R.M. and D.R.L. designed the study and analyzed the data; K.R.M., K.A.O., and A.D. performed mouse experiments. K.R.M. performed intravital multiphoton imaging. K.R.M and M.R.L. performed image analysis and quantification. C.N. and S.P.S. provided human biological samples and related quantitative analysis. K.R.M. and D.R.L. wrote the manuscript, with input from the other authors. D.R.L. supervised the research.

## Methods

### Mice

Mice were bred and maintained in the Alexandria Center for the Life Sciences animal facility of the New York University School of Medicine, in specific pathogen-free conditions. Albino B6 (B6(Cg)-*Tyr^c-2J^*/J, Jax 000058), Csf1r^EGFP^ (B6.Cg-Tg(Csf1r-EGFP)1Hume/J, Jax 018549), Ccr2^RFP^ (B6.129(Cg)-Ccr2tm2.1Ifc/J, Jax 017586), R26^mTmG^ (B6.129(Cg)-Gt(ROSA)26Sortm4(ACTB-tdTomato,-EGFP)Luo/J, Jax 007676), R26^nTnG^ (B6N.129S6-Gt(ROSA)26Sortm1(CAG-tdTomato*,-EGFP*)Ees/J, Jax 023537),

LysM^Cre^ (B6.129P2-Lyz2tm1(cre)Ifo/J, Jax 004781), Rac1^f/f^ (Rac1tm1Djk/J, Jax 005550), and CAG-dsRed (B6.Cg-Tg(CAG-DsRed*MST)1Nagy/J, JAX 006051) mice were purchased from Jackson Laboratories. *R26^dsRed^* mice were described previously (Luche et al., 2007) and were obtained from the laboratory of Dr. Gordon Fishell.

*Cx3cr1^CreER^*, *Cx3cr1^GFP^, and Cx3cr1^DTR^* were generated in our laboratory and have been described (Parkhurst, et al., 2013; Jung, et al., 2000; Diehl, et al., 2013). All experimental mice for this study were albino (homozygous for *Tyr^c-2J^*) as is required for intravital imaging in the skin. Cre-induction for the lineage tracing or total CAM labeling experiments was induced with a single intraperitoneal injection of Tamoxifen (Sigma-Aldrich; T5648) (100µg or 4mg in corn oil, respectively) in 1 month old mice. *Rac1^f/f^* recombination was induced with two intraperitoneal injections of Tamoxifen (2mg in corn oil) 48hrs apart in 1 month old mice. All imaging and experimental manipulations were performed on non-hairy mouse plantar (hind paw) skin. Preparation of skin for intravital imaging were performed as described below. Briefly, mice were anesthetized with intraperitoneal injection of ketamine/xylazine (15 mg/ml and 1 mg/ml, respectively in

PBS). After imaging, mice were returned to their housing facility. For subsequent revisits, the same mice were processed again with injectable anesthesia. The plantar epidermal regions were briefly cleaned with PBS pH 7.2, mounted on a custom-made stage, and a glass coverslip was placed directly against the skin. Anesthesia was maintained throughout the course of the experiment with vaporized isoflurane delivered by a nose cone. Mice from experimental and control groups were randomly selected for live imaging experiments. All lineage tracing and ablation experiments were repeated in at least three different mice. All animal procedures were performed in accordance with protocols approved by the Institutional Animal Care and Usage Committee of New York University School of Medicine.

### Intravital microscopy and laser ablation

Image stacks were acquired with an Olympus multiphoton FVMPE-RS system equipped with both InSight X3 and Mai Tai Deepsee (Spectra-Physics) tunable Ti:Sapphire lasers, using Fluoview software. For collection of serial optical sections, a laser beam (860nm for Hoechst 33342, 940nm for GFP/tdTomato/dsRed/RFP/Rhodamine/Second Harmonic Generation, 1200nm for Alexa Fluor 647, and 1300nm for Third Harmonic Generation, respectively) was focused through a water immersion lens (N.A. 1.05; Olympus) and scanned with a field of view of 0.5mm^2^, at 600 Hz. Z-stacks were acquired in 1-2μm steps for a ∼50-100μm range, covering the epidermis and dermis.

For all animal imaging, 1mm x 2 mm imaging fields (ROIs) were acquired with only the second harmonic signal (collagen) as a reference guide to the same anatomical position (1mm proximal of the most proximal walking pad on the mouse paw plantar skin.

Capillary blood flow was visualized in some experiments through intravenous injection with 18 mg/kg of dextran-rhodamine 70 kD (Sigma-Aldrich; R9379). Cell tracking analysis was performed by re-visiting the same area of the dermis in separate imaging experiments through using inherent landmarks of the skin to navigate back to the original region, including the distinct organization of the superficial vasculature networks. Cells that were unambiguously separated (by at least 250µm) from another were sampled to ensure the identity of individual lineages. For time-lapse recordings, serial optical sections were obtained between 5–10-minute intervals, depending on the experimental setup. Laser-induced cell ablation, capillary clot, or tissue damage was carried out with the same optics as used for acquisition. An 940nm laser beam was used to scan the target area (1-500μm^2^) and ablation was achieved using 50-70% laser power for ∼1sec. Ablation parameters were adjusted according to the depth of the target (10-50µm). Mice from experimental and control groups were randomly selected for live imaging experiments. All lineage tracing and ablation experiments were repeated in at least three different mice.

### In situ staining of neutrophils for intravital microscopy

Anesthetized mice were given fluorescently labeled antibodies via intravenous retroorbital injection immediately prior to imaging. Neutrophils were identified with 4μg of anti-Gr1-AF647 (BioLegend; clone RB6-8C5; Cat#108418).

### Drug treatments

To induce macrophage depletion, mice received intradermal injections of either Clodronate-liposomes or PBS-liposomes (stock concentration 5mg/ml; Liposoma; CP-005-005) (5µl per paw) every 3 days. Depending on experimental details, Cx3cr1^DTR^ mice received either intraperitoneal (IP) injection of diphtheria toxin (Sigma-Aldrich; D0564) every other day or a single low dose at 25ng/g body weight in PBS. To induce macrophage expansion, mice received daily intradermal injections of CSF1-FC (Bio-Rad; PPP031) or PBS in contralateral hind paws (5µl per paw) for 4 days.

### Generation of bone marrow chimeric reconstituted mice

Bone-marrow mononuclear cells were isolated from *Csf1r-GFP*;*CAG-dsRed* mice by flushing the long bones. Red blood cells were lysed with ACK lysing buffer and the remaining cells were resuspended in PBS for retroorbital injection. 4 × 10^6^ cells were then injected intravenously into 6-to 8-week-old *Csf1r-GFP* mice that were irradiated 4hr before reconstitution using 1,000 rads per mouse (2 × 500 rads, at an interval of 2hr, at X-RAD 320 X-Ray Irradiator). During irradiation, hind paws of recipient mice were lead-shielded to prevent any irradiation-induced loss of resident macrophage populations from our imaging area. At 1-and 10-months post irradiation, peripheral blood samples were collected from the submandibular (facial) vein in tubes containing EDTA (BD Biosciences; Dipotassium EDTA Microtainer; Cat# 365972). Red blood cells were lysed with ACK lysing buffer and the remaining cells were subjected flow cytometric analysis on a LSR II with FACSDiva and FlowJo 10.10.1 software (BD Biosciences) to check for reconstitution (Extended Data Fig. 10b).

### Skin whole mount staining

Whole skin was collected from the hind paw and fixed in 4% paraformaldehyde in PBS overnight at 4°C, washed in PBS, permeabilized and blocked for 1hr (2% Triton-X, 5% normal donkey serum and 1% BSA in PBS). For CD206 staining, blocked tissue was incubated in Alexa Fluor 647 rat anti-CD206 (1:500; Biolegend C068C2) overnight at 4°C, washed in PBS with 2% Triton-X, washed with PBS, and then mounted on a slide with Prolong Gold antifade mounting medium (Invitrogen) with a #1.5 coverslip. Nuclear counterstaining was achieved by performing a single intravenous injection of Hoechst 33342 (15 mg/kg) 30 minutes prior to mouse euthanasia. Whole mount skin samples were imaged with the same imaging conditions and setup used for intravital microscopy.

### Human skin samples

Written informed consent was obtained for postmortem examination from next of kin for all patients. Clinical information and laboratory data were obtained from the electronic medical record. Sex and gender information was not used. Patients in young group were below 40 years of age. Patients in old group were above 40 years of age. Patients with skin or vascular pathologies were excluded. Skin samples were obtained from the anterolateral chest and fixed in 10% formalin for at least 24h prior to processing. Slides were stained with hematoxylin and eosin, CD68 (Clone 514H12) and ERG (Clone EPR3864). Macrophages and capillaries were identified using a combination of morphology, CD68 and ERG staining. The age group of samples were blinded to the researcher and counting was performed on at least eight high-power fields (40x) within 100um of the epidermis.

### Image Analysis

Raw image stacks were imported into Fiji (NIH, USA) or Imaris software (Bitplane/Perkin Elmer) for further analysis. Provided images and supplementary videos are typically presented as a maximal projection of 4-8µm optical sections. For visualizing individual labeled cells expressing the dsRed or tdTomato Cre reporters, the brightness and contrast were adjusted accordingly for the green (GFP) and red (dsRed/tdTomato) channels and composite serial image sequences were assembled as previously described. Images were obtained as large, tiled image stacks at roughly the same positions and then manually aligned over the experimental time course in Imaris (Bitplane/Perkin Elmer) by using data from all channels. Random ROI were selected for image analysis. Quantification of CAM coverage and capillary blood flow was performed blinded to minimize researcher bias.

### Statistical Analysis

Data are expressed as mean ± SD. An unpaired Student’s *t*-test was used to analyze data sets with two groups. One-way ANOVA was used to analyze data sets with three or more groups. *p* < 0.05 to *p* < 0.0001 indicated a significant difference. Statistical calculations were performed using the Prism software package (GraphPad, USA).

---

**Extended Data Figure 1.**
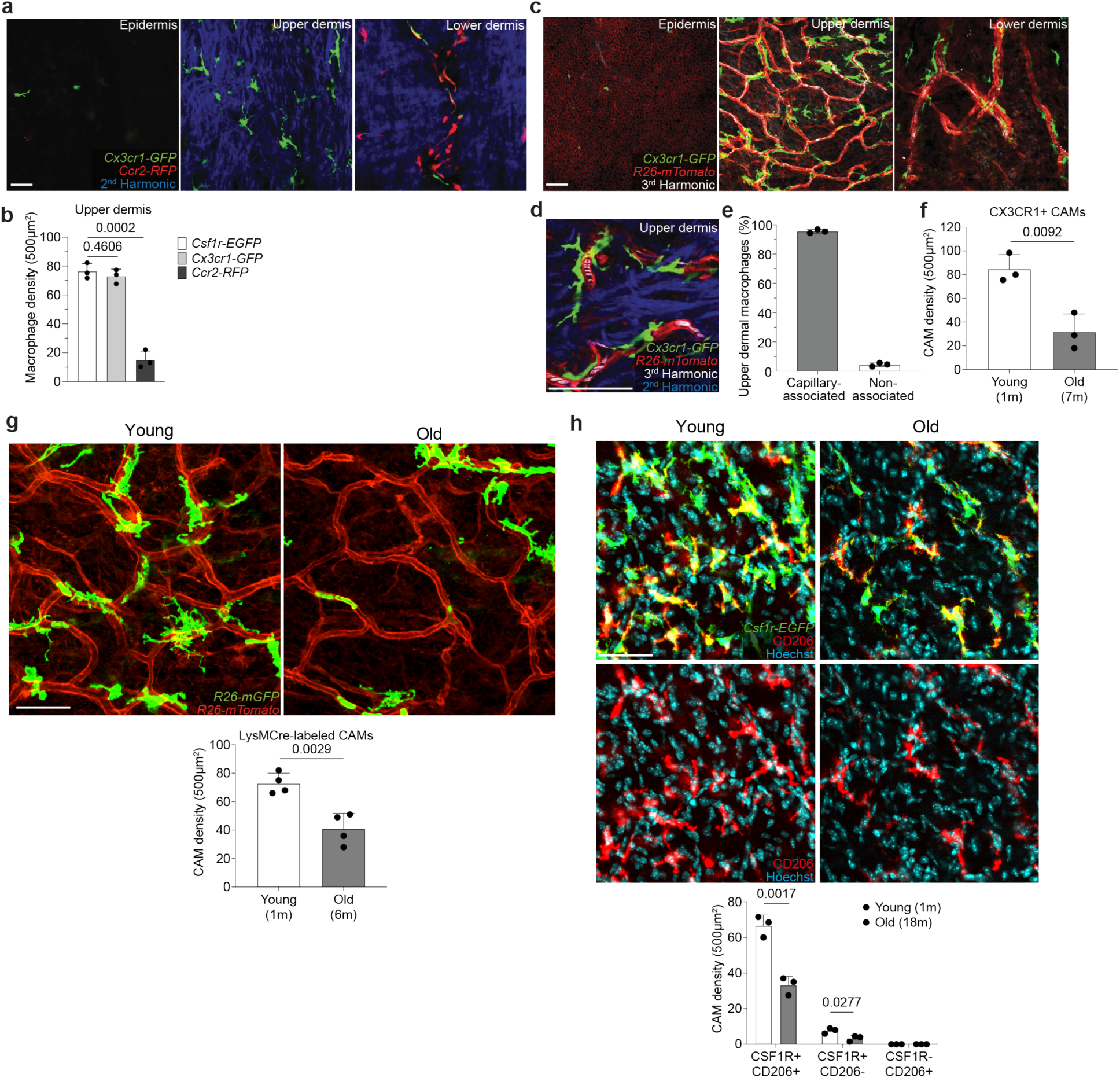
Upper dermal macrophages are capillary-associated and decline in density with age. **a,** Representative images of cells expressing *Ccr2-RFP*, *Csf1r-EGFP*, or *Cx3cr1-GFP* in the epidermis, upper dermis, and lower dermis of 1 month old mice. **b,** Quantifications of cells expressing *Csf1r-EGFP*, *Cx3cr1-GFP, or Ccr2-RFP*, in the upper dermis (n = 3 mice in each group; two 500µm^2^ regions per mouse; Cell number (*Csf1r-EGFP* vs *Cx3cr1-GFP; Csf1r-EGFP* vs *Ccr2-RFP*) was compared by unpaired Student’s t test; mean ± SD). **c,** Visualization of CX3CR1-expressing cells in all skin layers: epidermis, upper dermis, and lower dermis (*Cx3cr1-GFP;R26-mTmG*) was performed during homeostatic conditions. **d,** Representative image of labeled upper dermal macrophages in contact with capillary superficial plexus (red) in 1 month old mice. **e,** Quantifications reveal most upper dermal macrophages are capillary-associated macrophages (CAMs) (n = 3 mice in total; three 500µm^2^ regions per mouse; mean ± SD). **f,** Quantifications of cells expressing *Cx3cr1-GFP* in the upper dermis (n = 3 mice in each group; two 500µm^2^ regions per mouse; CAM density (1 vs 7-month-old) was compared by unpaired Student’s t test; mean ± SD). **g,** Representative optical sections of CAMs in young (1-month-old) and old (6-month-old) *LysM-Cre;R26-mTmG* mice. Quantifications of membrane-GFP+ cells in the upper dermis (n = 4 mice in each group; two 500µm^2^ regions per mouse; CAM density (1 vs 6-month-old) was compared by unpaired Student’s t test; mean ± SD). **h,** Representative upper dermal optical sections from whole mount skin samples of young (1-month-old) and old (18-month-old) *Csf1r-EGFP* mice stained with AF647 anti-mouse CD206 (clone C068C2) antibody. Quantifications for CD206 and CSF1R co-expression in the upper dermis (n = 3 mice in each group; two 500µm^2^ regions per mouse; CAM density (1 vs 18-month-old) was compared by unpaired Student’s t test; mean ± SD). Scale bar, 50 µm.

**Extended Data Figure 2.**
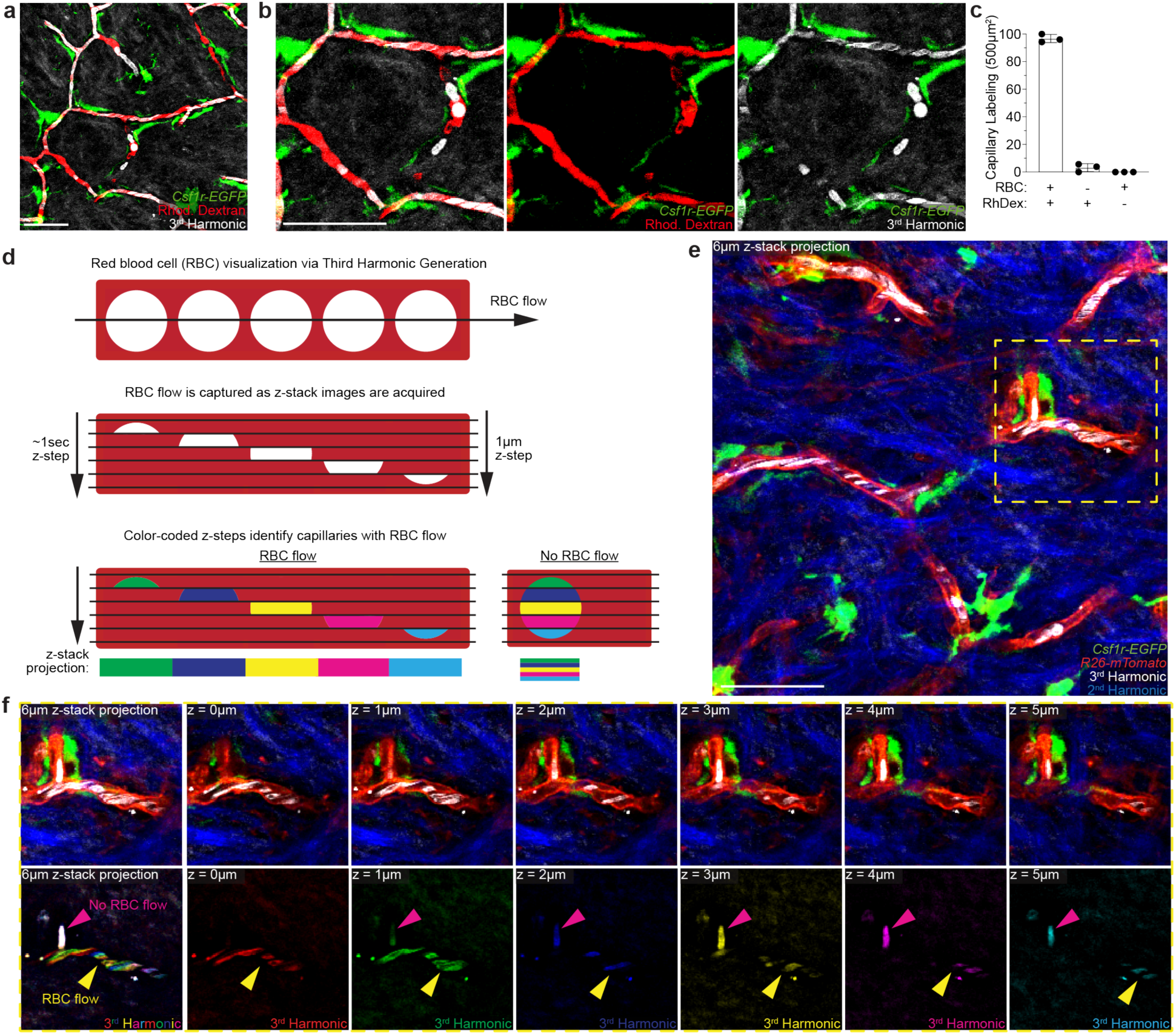
Label-free *in vivo* visualization of capillary blood flow via third harmonic effect generated by red blood cells. **a-b**, Simultaneous visualization of blood flow through the superficial capillary plexus via third harmonic generation (white) from red blood cells (RBC) and intravenous rhodamine dextran (RhDex) (red). **c**, Quantifications of RBC and RhDex labeling of the upper dermal superficial capillary plexus (n = 3 mice in total; four 500µm^2^ regions per mouse; mean ± SD). **d,** Scheme of red blood cell (RBC) flow through a segment of the superficial skin capillary network. During three-dimensional image acquisition, flowing red blood cells are captured at different x,y positions for each z-section along the capillary segment. Pseudo coloring each z-step through a capillary segment distinguishes flowing RBCs as a multicolor patchwork or rainbow-effect and leaves obstructed/non-flowing RBCs as white (full overlap of all colors). **e,** Representative image of RBC visualization in the upper dermis in *Csf1r-EGFP; R26-mTmG* mice. **f,** Representative optical z-sections through upper dermal capillaries. Third harmonic signal is pseudo colored differently for each z-step to visualize RBC movement along the capillary segments during image acquisition. Scale 50µm.

**Extended Data Figure 3.**
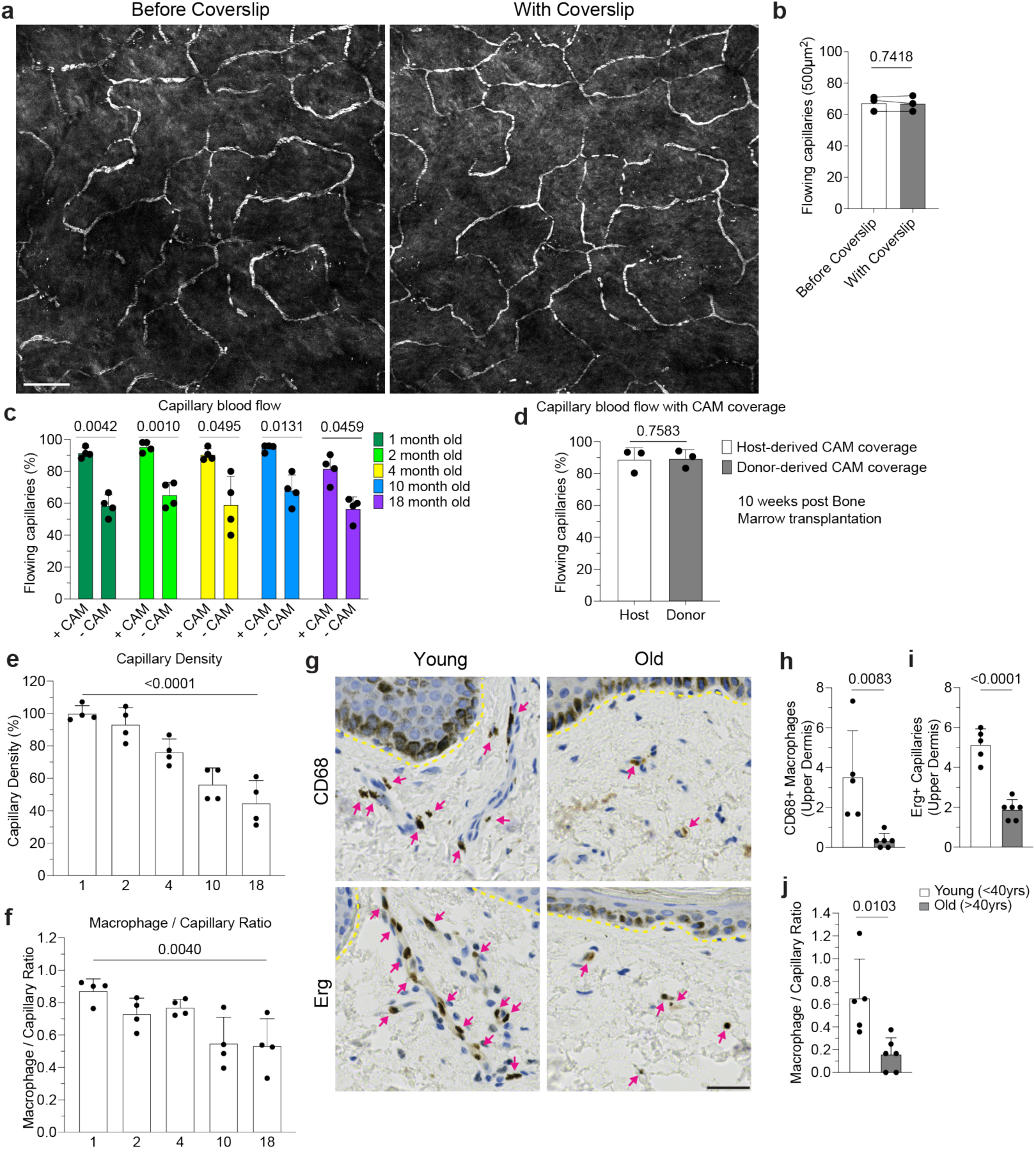
Capillary flow and associated macrophage loss in mice and humans. **a,** Representative images of capillary blood flow in WT mice before and after placement of a coverslip on top of hind paw skin of 6 month old mice. **b,** Quantification of capillary blood flow (n = 3 mice in total; two 500µm^2^ regions per mouse; capillary blood flow (Before Coverslip vs With Coverslip) was compared in the same imaging areas by paired Student’s t test; mean ± SD). **c,** Quantification of capillaries with blood flow as measured by stalled RBCs as described in Extended Data Figure 2 in 1, 2-, 4-, 10-, and 18-month-old mice (n = 878 CAM+ capillary segments, n = 215 CAM-capillary segments; n = 4 mice in each age group; capillary blood flow (CAM+ vs CAM-) was compared by paired Student’s t test; mean ± SD). **d,** Quantification of capillaries with blood flow in in bone marrow chimeras with *Csf1r-GFP*;*CAG-dsRed* bone marrow transferred into lethally irradiated *Csf1r-GFP* mice (n = 207 host-derived CAM+ capillary segments, n = 79 donor-derived CAM+ capillary segments; n = 3 mice in total; CAM+ capillary blood flow (Host-derived vs Donor-derived) was compared by paired Student’s t test; mean ± SD). **e,f** Quantification of capillary niche age-associated changes, (e) Capillary density and (f) CAM / Capillary segment ratio (n = 4 mice in each age group; two 500µm^2^ regions per mouse; comparison across age groups was by one-way ANOVA; mean ± SD). **g,** Representative immunohistochemistry of CD68+ macrophage and Erg+ capillary density in the upper dermis of both young (<40y) and old (>40y) human samples. **h-j,** Quantification of capillary niche age-associated changes: (h) CD68+ macrophage density, (i) Erg+ capillary density, (j) Upper dermal macrophage / Capillary endothelium ratio (n = 5 patient samples in each group; three imaging regions per sample; Comparison by unpaired Student’s t test; mean ± SD). Scale bar, 50µm.

**Extended Data Figure 4.**
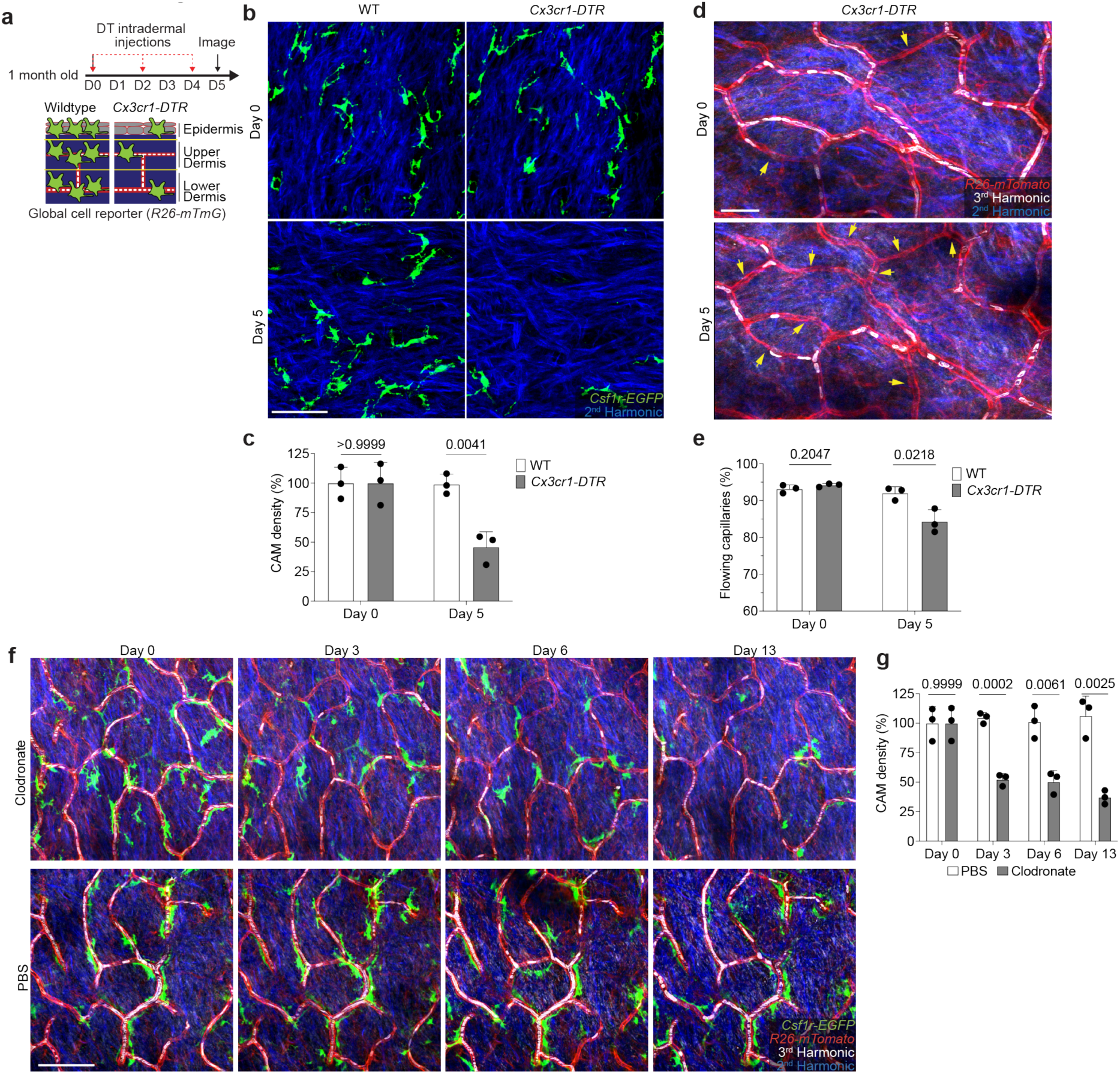
Impaired skin capillary blood flow following acute macrophage depletion. **a,** Scheme of capillary blood flow tracking following intraperitoneal injection every other day with diphtheria toxin (DT) (25ng/g body weight in PBS) in both *Cx3cr1-DTR*; *R26-mTmG* and WT control (*R26-mTmG*) mice. **b,** Representative revisits of the same upper dermal capillary niches to visualize CAMs (*Csf1r-EGFP*) and dermal collagen (Second Harmonic). **c,** Quantifications reveal a significant reduction in CAM density following DT-treatment (n = 3 mice in each group; two 500µm^2^ regions per mouse; CAM density (WT vs *Cx3cr1-DTR*) was compared for Day 0 and Day 5 time points by unpaired Student’s t test; mean ± SD). **d,** Representative revisits of the same capillary network to visualize capillaries (*R26-mTmG*) and RBC flow (Third Harmonic). **e,** Quantifications reveal a significant reduction in the percentage of capillaries with blood flow following DT-induced cell depletion (n = 3 mice in each group; two 500µm^2^ regions per mouse; obstructed capillary flow (WT vs *Cx3cr1-DTR*) was compared for Day 0 and Day 5 time points by unpaired Student’s t test; mean ± SD). **f,** Representative images demonstrate macrophage depletion following intradermal injections of clodronate-liposomes every 3 days. Repeated intravital imaging of the vascular niche was performed to visualize macrophages (*Csf1r-GFP*), capillaries (*R26-mTmG*) and RBC flow (Third Harmonic). **g,** Quantification of CAM density following macrophage depletion (n = 3 mice in each group; two 500µm^2^ regions per mouse; CAM density (clodronate vs PBS) was compared by unpaired Student’s t test; mean ± SD). Scale bar, 50 µm.

**Extended Data Figure 5.**
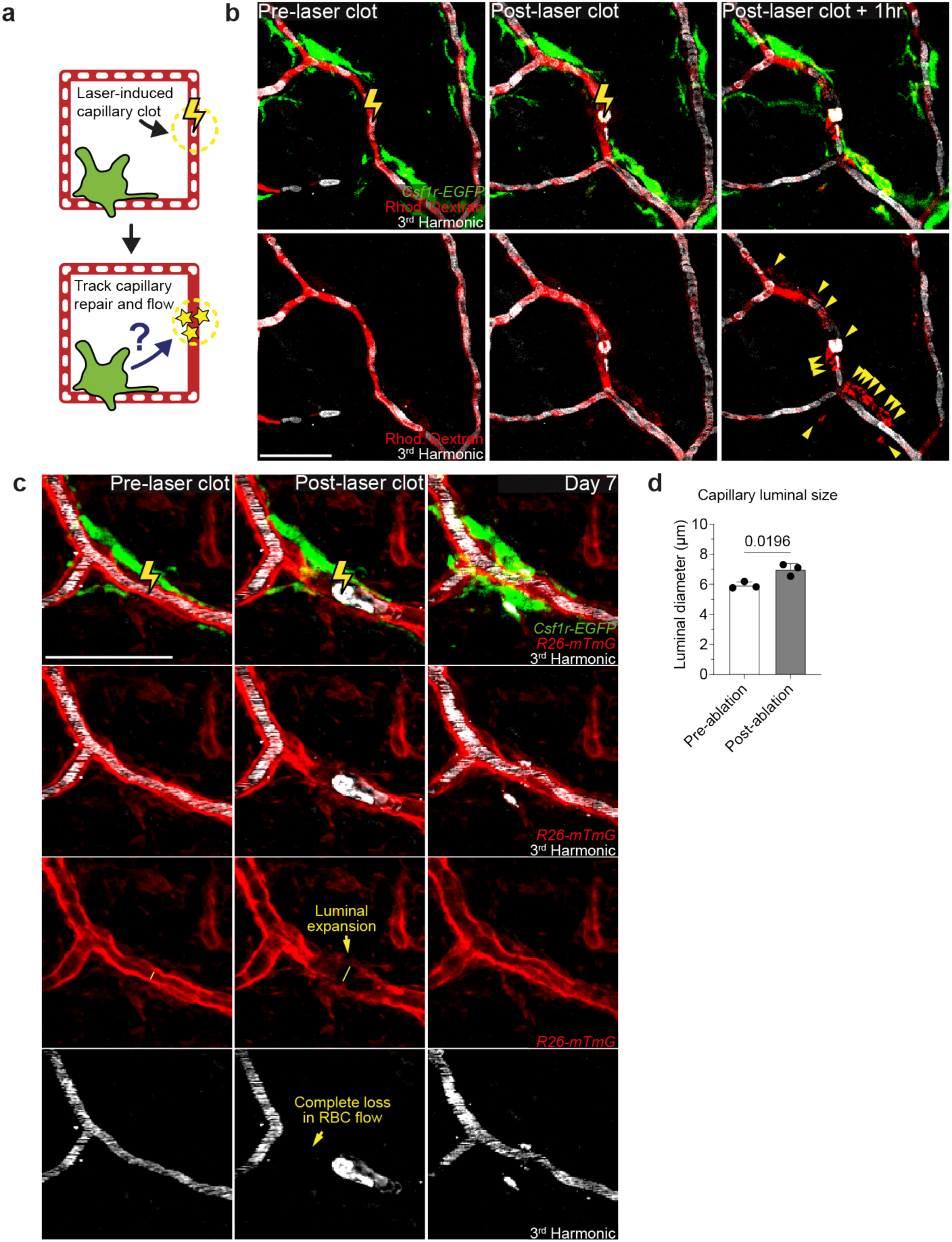
Laser-induced model of acute capillary clot formation and repair. **a,** Scheme of laser-induced capillary clot experiment. **b,** Sequential revisits of damaged capillary niche after laser-induced clot formation in *Csf1r-EGFP* mice. Simultaneous visualization of blood flow before and after clot formation via third harmonic generation (white) from red blood cells (RBC) and intravenous rhodamine dextran (RhDex) (red). Yellow lightning bolt indicates site of laser-induced capillary clot. Yellow arrowheads indicate extra-luminal vascular debris. **c,** Sequential revisits of damaged capillary niche after laser-induced clot formation in *Csf1r-EGFP;R26-mTmG* mice. Capillary clot formation (yellow lightning bolt) was performed at 940nm for 1s in 4-month-old mice. **d,** Quantification of capillary lumen diameter before and immediately after laser-induced clotting (n = 24 capillary clots; 3 mice in total; capillary luminal diameter (pre-ablation vs post-ablation) was compared by paired Student’s t test; mean ± SD). Scale bar, 50 µm.

**Extended Data Figure 6.**
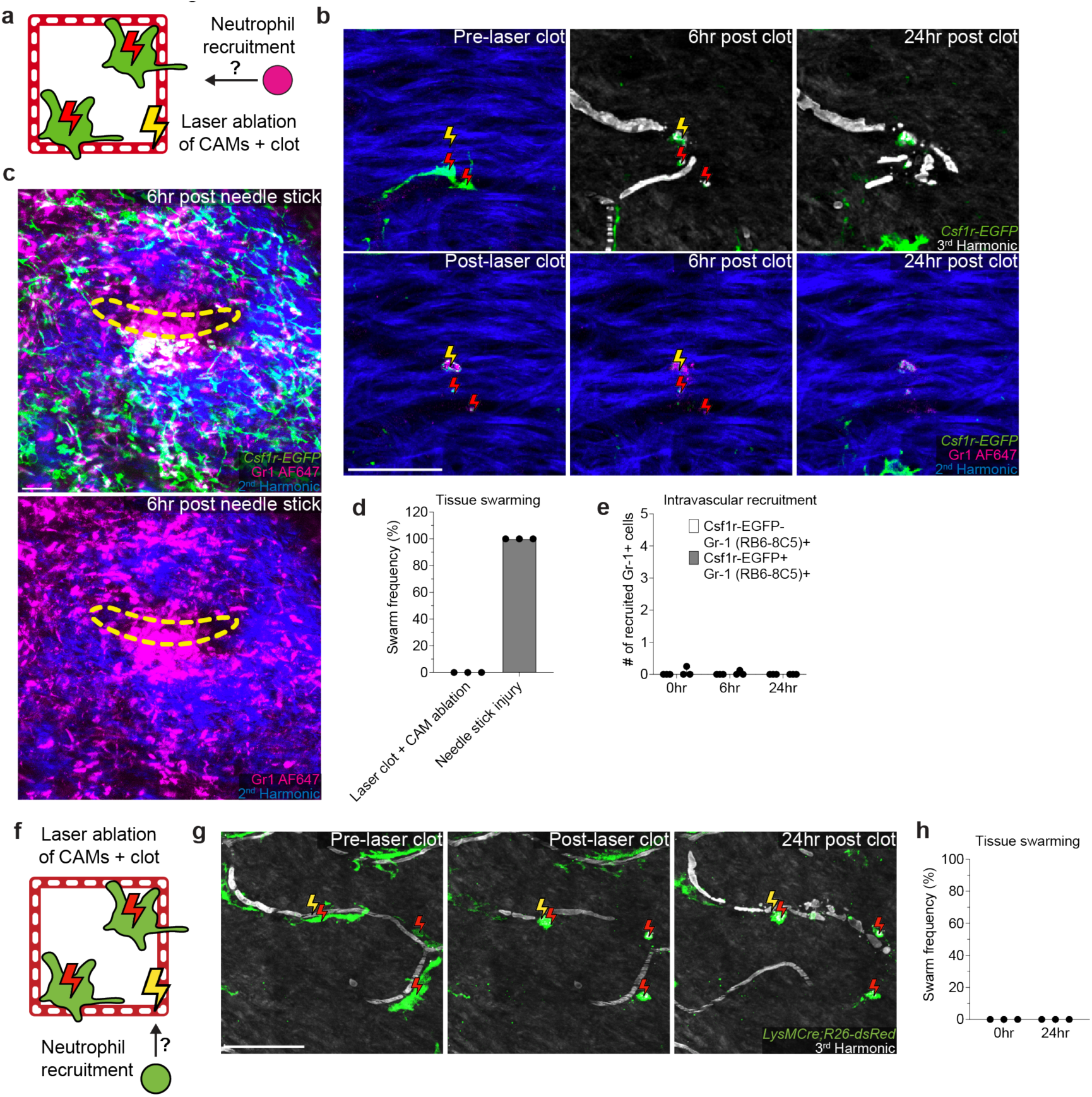
Laser-induced clots and CAM ablations do not recruit neutrophil swarms. **a,** Scheme of tracking neutrophil recruitment and swarming after laser-induced capillary clot experiment. **b-c,** (b) Sequential revisits of damaged capillary niche after laser-induced CAM ablation and clot formation or (c) large non-sterile tissue damage (28-gauge needle stick) in *Csf1r-EGFP* mice with intravenous neutrophil antibody labeling (Gr-1, clone RB6-8C5). Macrophage laser ablation (red lightning bolt) and capillary clot formation (yellow lightning bolt) were both performed at 940nm for 1s. Needle injury (yellow dashed line) was performed through epidermis and dermis. **d,** Quantification of neutrophil swarming at 6 hours tissue damage (n = 16 capillary clots in CAM ablated regions, n = 9 needle injuries; 3 mice in each group; mean ± SD). **e,** Quantification of intravascular recruitment of neutrophil and monocyte populations at 0, 6, and 24 hours post laser-induced capillary clotting in CAM ablated regions (n = 16 capillary clots in CAM ablated regions, 3 mice in total; mean ± SD). **f,** Scheme of tracking neutrophil recruitment and swarming after laser-induced capillary clot experiment. **g,** Sequential revisits of damaged capillary niche after laser-induced CAM ablation and clot formation in *LysMCre;R26-dsRed* mice. Macrophage laser ablation (red lightning bolt) and capillary clot formation (yellow lightning bolt) were both performed at 940nm for 1s. **h,** Quantification of neutrophil swarming at 6 hours tissue damage (n = 32 capillary clots in CAM ablated regions; 3 mice in total; mean ± SD). Scale bar, 50 µm.

**Extended Data Figure 7.**
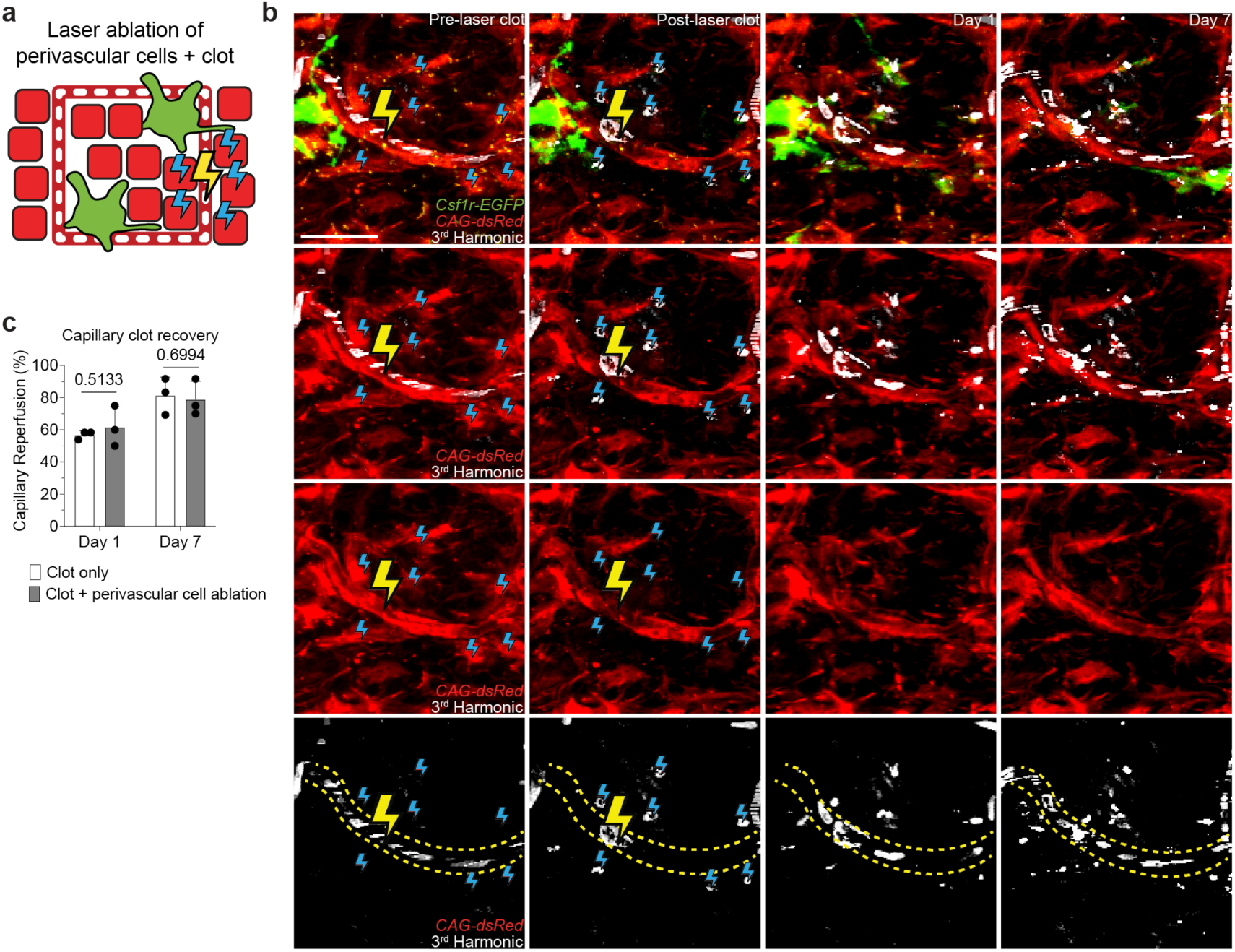
Laser ablation of non-macrophage perivascular dermal cells does not impair capillary clot repair and reperfusion. **a,** Scheme of laser-induced capillary clot experiment. **b,** Sequential revisits of damaged capillary niche after laser-induced perivascular dermal cell ablation and clot formation in *Csf1r-EGFP*;*CAG-dsRed* mice. Perivascular dermal cell laser ablation (blue lightning bolt) and capillary clot formation (yellow lightning bolt) were both performed at 940nm for 1s. **c,** Quantification of capillary reperfusion at Day 1 and 7 after laser-induced clotting and perivascular cell ablation (n = 34 capillary clots in CAM ablated regions, n = 37 capillary clots in control regions; 3 mice in total; capillary reperfusion (perivascular dermal cell ablation vs control) was compared by paired Student’s t test; mean ± SD). Scale bar, 50 µm.

**Extended Data Figure 8.**
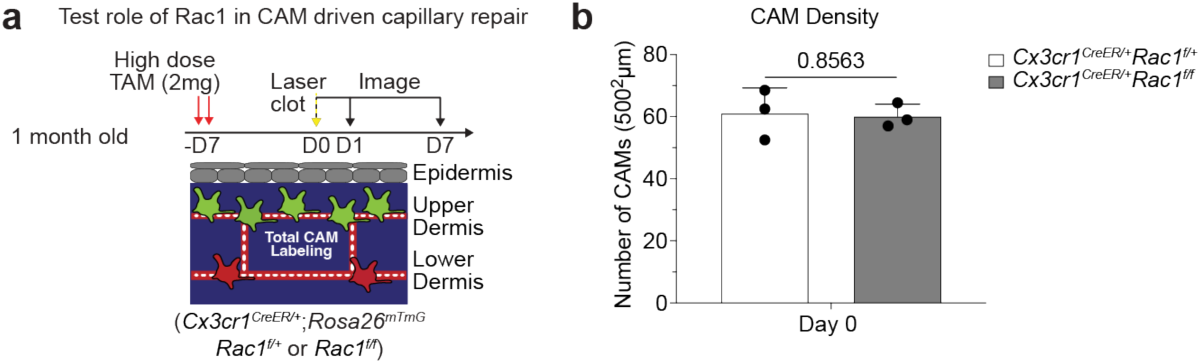
CAM density in mice with deficiencies in macrophage phagocytic machinery. **a,** Scheme of laser-induced capillary clot in *Cx3cr1^CreER^*;*Rac1^fl/fl^* and *Cx3cr1^CreER^*;*Rac1^fl/+^* mice. **b,** Quantification of CAM density at Day 7 after laser-induced clotting (n = 3 mice in each group; two 500µm^2^ regions per mouse; CAM density (*Rac1^fl/+^* vs *Rac1^fl/fl^*) was compared at day 0 (same day of clot induction) by unpaired Student’s t test; mean ± SD).

**Extended Data Figure 9.**
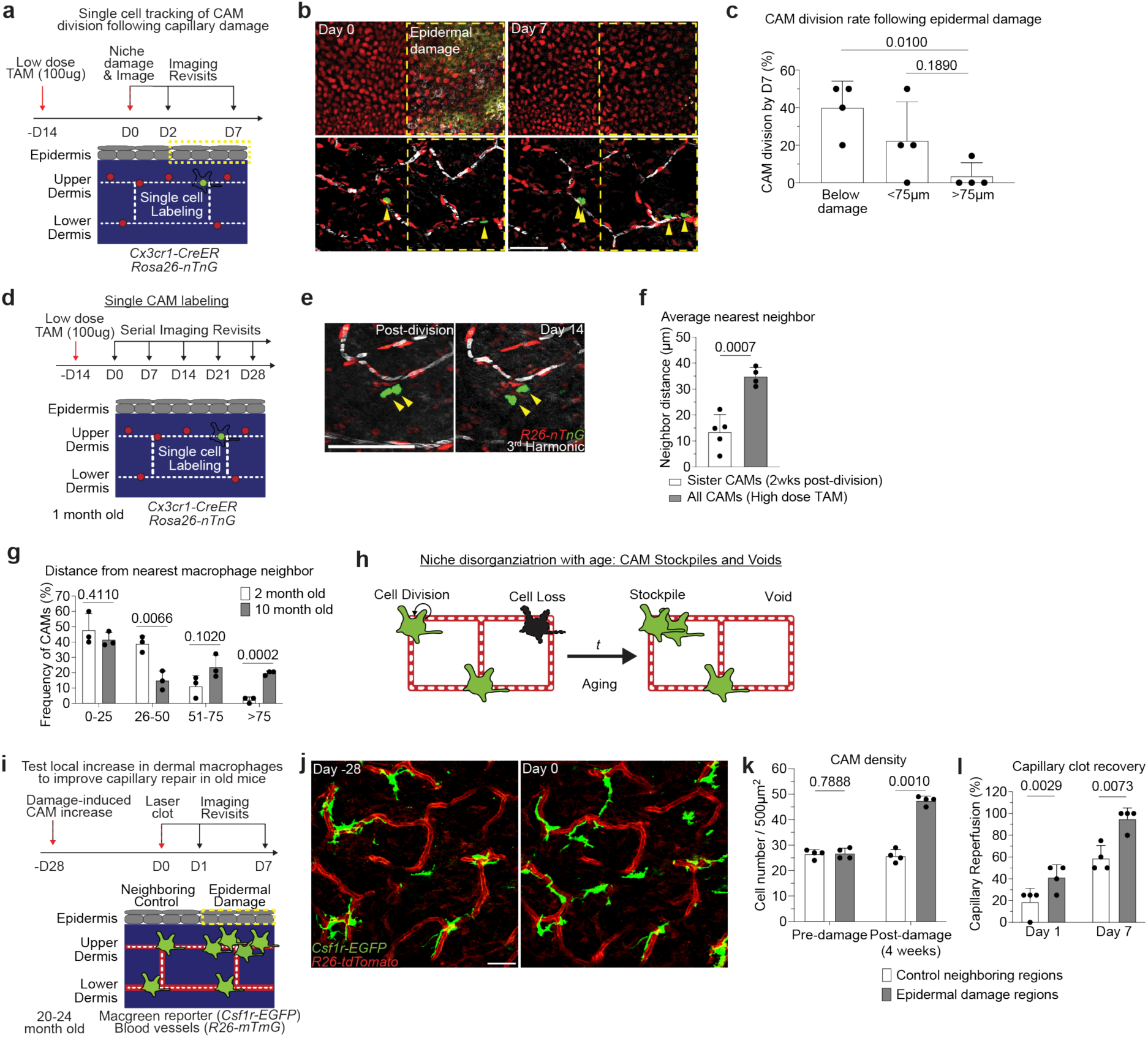
CAM loss and disorganization is partially restored by local epidermal damage to improve capillary repair in old mice. **a,** Scheme of tracking of CAM proliferation following laser-induced damage to nearby capillary or epidermal niches. **b,** Representative revisits of single macrophage lineage tracing in *Cx3cr1-CreERT2*; *R26-nTnG* mice following epidermal damage. **c,** Quantification of CAM proliferation based on proximity to epidermal damage (n = 25 CAMs tracked in capillary damage regions, n = 56 CAMs tracked in epidermal damage regions; 4 mice in each group; CAM proliferation (based on damage proximity) was compared at day 7 by paired Student’s t test; mean ± SD). **d,** Scheme of long-term tracking of CAM migration following cell division. **e,** Representative revisits of single macrophage lineage tracing in *Cx3cr1-CreERT2*; *R26-nTnG* mice. Weekly revisits were performed during homeostatic conditions for 5 weeks following a single low-dose intraperitoneal injection of tamoxifen (50µg). **f,** Quantification of neighboring CAM distance of recently divided sister CAMs was compared to total CAM neighboring distance from *Cx3cr1-CreERT2*; *R26-nTnG* mice given a single high-dose intraperitoneal injection of tamoxifen (4mg) (n = 26 sister CAM pairs, from 5 mice in low-dose group; n = 188 CAMs, from 4 mice in high-dose group; Average nearest neighbor distance (2 vs 10 month old) was compared at 0-25, 26-50, 51-75, and >75µm intervals by unpaired Student’s t test mean ± SD). **g,** Quantification of distance between nearest CAM neighbors in 2-and 10-month-old *Csf1r-EGFP* mice (n = 90 CAMs in 2-month-old mice, n = 101 CAMs in 10-month-old mice; 3 mice in each age group; Frequency of CAM distribution (2 vs 10 month old) was compared at 0-25, 26-50, 51-75, and >75µm intervals by unpaired Student’s t test; mean ± SD). **h,** Working model of macrophage renewal and organization in the skin aging capillary niche. **i,** Scheme of local damage-induced expansion of CAMs in the aged capillary niche. **j,** Representative images of CAM density in *Csf1r-EGFP; R26-mTmG* mice 28 days following overlaying epidermal laser damage. **k,** Quantification of CAM density following overlaying epidermal damage (n = 4 mice in total; two 500µm^2^ regions for both epidermal damage and control conditions per mouse; CAM density (Epidermal damage vs Control) was compared by paired Student’s t test; mean ± SD). **l,** Quantification of capillary reperfusion at day 1 and 7 after laser-induced clotting (n = 17 capillary clots in epidermal damage regions, n = 17 capillary clots in neighboring control regions; 4 mice in total; capillary reperfusion (with epidermal damage vs control) was compared at day 1 and day 7 by paired Student’s t test; mean ± SD). following laser-induced damage to overlaying epidermal niche. Scale bar, 50µm.

**Extended Data Figure 10.**
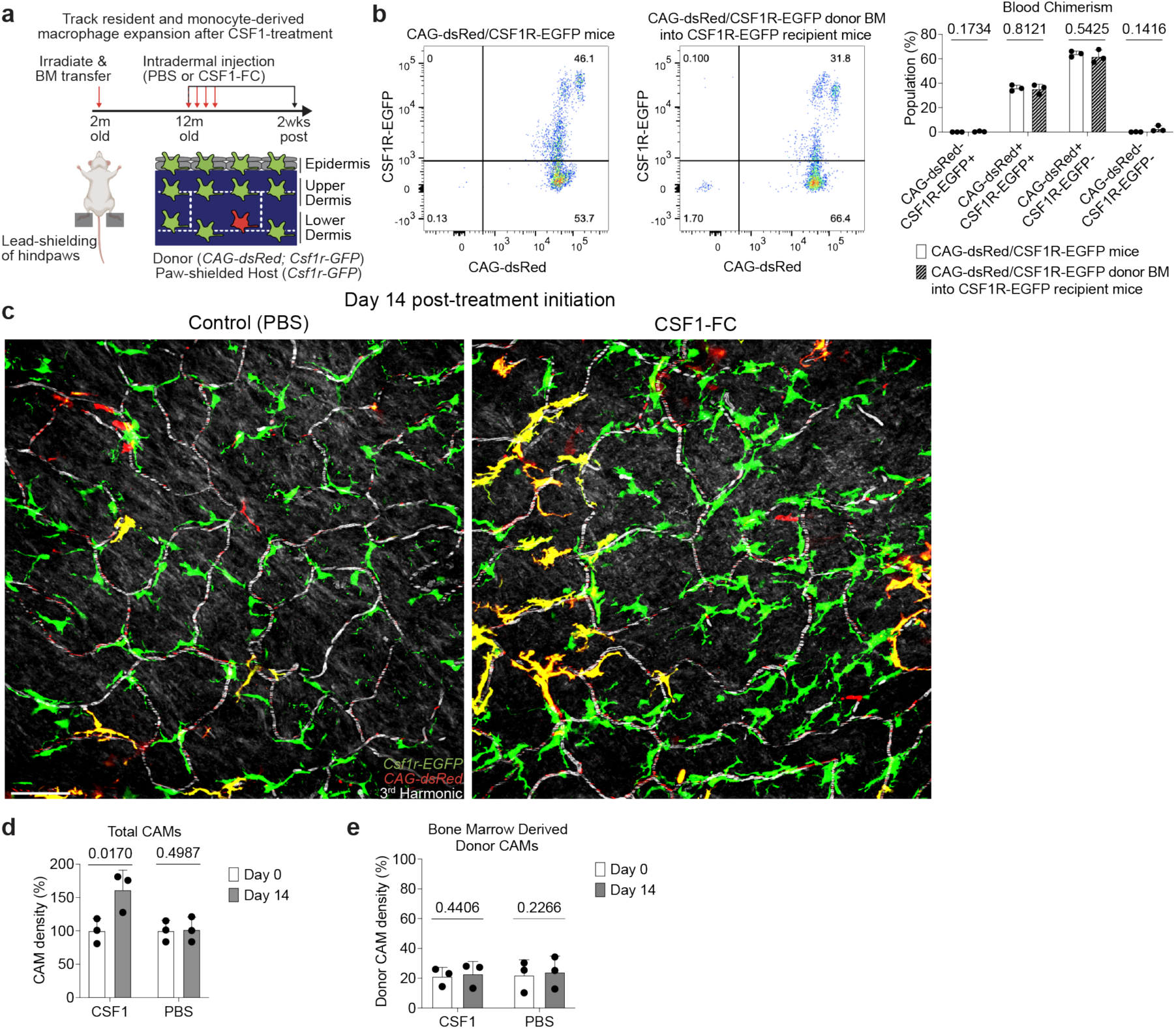
Intradermal CSF1 treatment locally expands CAM populations in old mice. **a,** Scheme of serial imaging of skin resident macrophage repopulation in bone marrow chimeras with *Csf1r-GFP*;*CAG-dsRed* bone marrow transferred into lethally irradiated *Csf1r-GFP* mice following CSF1-treatment. Note hind paws of these mice were lead-shielded to prevent irradiation-induced loss of resident macrophage populations from our imaging area. **b,** Quantification of blood chimerism 10 months after *Csf1r-GFP*;*CAG-dsRed* bone marrow transfer into lethally irradiated *Csf1r-GFP* mice (n = 3 mice in each group; Percentage of cells from blood (*Csf1r-GFP*;*CAG-dsRed* mice vs bone marrow transferred mice) was compared for dsRed-/GFP+, dsRed+/GFP+, dsRed+/GFP-, dsRed-/GFP-populations by unpaired Student’s t test; mean ± SD). **c,** Representative images of CAM density 14 days following daily intradermal injections (4 days) of CSF1-Fc (porcine CSF1 and IgG1a Fc region fusion protein) and PBS in contralateral hind paws of 12-month-old mice. **d,** Quantification of capillary-associated macrophage density following CSF1-Fc or PBS treatment (n = 3 mice in total; two 500µm^2^ regions of each treatment condition per mouse; Percentage of CAM density change (Day 0 vs 14) was compared for CSF1-Fc and PBS regions by paired Student’s t test; mean ± SD). **e,** Quantification of donor bone marrow-derived (GFP+/dsRed+) CAM density following CSF1-Fc or PBS treatment (n = 3 mice in total; two 500µm^2^ regions of each treatment condition per mouse; Percentage of donor bone marrow-derived CAM density change (Day 0 vs 14) was compared for CSF1-Fc and PBS regions by paired Student’s t test; mean ± SD). Scale bar, 50µm.

## Supplemental Video Legends

**Supplemental Video 1.** Serial optical sections through mouse plantar skin, including epidermis (0-25µm), upper dermis (26-50µm), and lower dermis (51-100µm) in *R26-mTmG* (red) mice. Note Second and Third Harmonic Generation illuminates dermal collagen (blue) and red blood cells (white), respectively. Scale bar, 50µm.

**Supplemental Video 2.** Serial optical sections through mouse plantar skin, including epidermis (0-15µm), upper dermis (16-40µm), and lower dermis (41-52µm) in *Csf1r-EGFP* mice. Note distinct macrophage (green) density and morphology in each tissue niche. Scale bar, 50µm.

**Supplemental Video 3.** Time-lapse recording of capillary blood flow via third harmonic generation (white) from red blood cells and intravenous rhodamine dextran (red) in *Csf1r-EGFP* mice. Note some capillary segments show obstructed red blood cell and rhodamine dextran flow. Scale bar, 50µm.

**Supplemental Video 4.** Serial optical sections through upper dermal capillary niche in *Cx3cr1-GFP; R26-mTmG* mice. Note macrophage (green) localization and morphology around capillary (red) network. Scale bar, 50µm.

**Supplemental Video 5.** Time-lapse recording of capillary blood flow in *Cx3cr1-CreER; R26-mTmG* mice. Note inconsistent and obstructed blood flow (white) in capillary (red) segments without associated macrophages (green). Scale bar, 50µm.

**Supplemental Video 6.** Time-lapse recording following laser-induced capillary clot formation in *Csf1r-GFP; R26-mTmG* mice. Note rapid migration of neighboring capillary-associated macrophages (green) toward site of clot formation (yellow arrowhead). Scale bar, 50µm.

## Notes

### Competing Interest Statement

D.R.L consults for and has equity interest in Vedanta Bioscience, Immunai, IMIDomics, Evommune, and Pfizer Pharmaceuticals.

### Summary of Updates

We are pleased to submit a revised version of our manuscript. We have performed new experiments and significantly revised the text, which has allowed us to strengthen our conclusions and to better highlight the novelty of our approach and findings.

